# Plasmin improves oedematous blood-gas barrier by cleaving epithelial sodium channels

**DOI:** 10.1101/2020.02.09.940619

**Authors:** Runzhen Zhao, Gibran Ali, Hong-Guang Nie, Yongchang Chang, Deepa Bhattarai, Xuefeng Su, Xiaoli Zhao, Michael A. Matthay, Hong-Long Ji

## Abstract

**Background and Purpose:** Lung oedema in association with suppressed fibrinolysis is a hallmark of lung injury. We aimed to test whether plasmin cleaves epithelial sodium channels (ENaC) to resolve lung oedema fluid.

**Experimental Approaches:** Human lungs and airway acid-instilled mice were used for analysing fluid resolution. *In silico* prediction, mutagenesis, *Xenopus* oocytes, immunoblotting, voltage clamp, mass spectrometry, protein docking, and alveolar fluid clearance were combined for identifying plasmin specific cleavage sites and benefits.

**Key Results:** Plasmin led to a marked increment in lung fluid resolution in both human lungs ex vivo and injured mice. Plasmin specifically activated αβγENaC channels in oocytes in a time-dependent manner. Deletion of four consensus proteolysis tracts (αΔ432-444, γΔ131-138, γΔ178-193, and γΔ410-422) eliminated plasmin-induced activation significantly. Further, immunoblotting assays identified 7 cleavage sites (K126, R135, K136, R153, K168, R178, K179) for plasmin to trim both furin-cleaved C-terminal fragments and full-length human γENaC proteins. In addition to confirming the 7 cleavage sites, 9 new sites (R122, R137, R138, K150, K170, R172, R180, K181, K189) in synthesized peptides were found to be cleaved by plasmin with mass spectrometry. These cleavage sites were located in the finger and the thumb, particularly the GRIP domain of human ENaC 3D model composed of two proteolytic centres for plasmin. Novel uncleaved sites beyond the GRIP domain in both α and γ subunits were identified to interrupt the plasmin cleavage-induced conformational change in ENaC channel complexes. Additionally, plasmin could regulate ENaC activity via the G protein signal.

**Conclusion and Implications:** We demonstrate that plasmin could cleave ENaC to benefit the blood-gas exchange by resolving oedema fluid as a potent fibrinolytic therapy for oedematous pulmonary diseases.

**Bullet point summary:** *What is already know:*

- Serine proteases proteolytically cleave epithelial sodium channels, including plasmin and uPA acutely.
- Activity of epithelial sodium channels is increased post proteolysis.

*What this study adds:*

- Plasmin cleaves up to 16 sites composed of two proteolytic centres in both full-length and furin-cleaved human γ subunit of epithelial sodium channels in hours.
- Non-proteolytic sites in both α and γ subunits interrupt the plasmin cleavage-induced channel gating.
- Intratracheally instilled plasmin facilitates alveolar fluid clearance in normal human and injured mouse lungs.

*Clinical significance:*

- Activation of human lung epithelial sodium channels by plasmin may benefit lung oedema resolution as a novel therapy for ARDS.

## 1. INTRODUCTION

One of the most common complications of sepsis is the acute respiratory distress syndrome (ARDS) with a hallmark of dysfunctional blood-gas barrier, occurring in nearly 200,000 patients per year with a mortality above 30% (Ware & Matthay, 2000). ARDS is characterized by hypoxemia with bilateral infiltrates of the lungs with noncardiac origin (Ranieri et al., 2012; Sartori & Matthay, 2002). Prolonged suppression of normal fibrinolytic activity, paralleled by a sustained increase in plasminogen activator inhibitor-1 (PAI-1) has been consistently observed in clinical and preclinical studies of septic ARDS (Bertozzi et al., 1990). Fibrinolytic activity is further impaired following the development of sepsis with multiorgan failure (Asakura et al., 2001). An elevated plasma PAI-1 concentration is associated with an adverse outcome in sepsis (Hermans et al., 1999). Caged plasminogen activators lose their ability to produce plasmin, a key molecule of the plasminogen/plasmin system that sustains normal fibrinolysis. The reduced fibrinolytic activity was also reported in bronchoalveolar fluid and pleural effusion of septic illnesses (Idell, James & Coalson, 1992). Concurrently, increased permeability pulmonary oedema often develops. Accumulation of alveolar oedema fluid mainly results from an increase in lung endothelial and epithelial permeability that cannot be compensated by transepithelial fluid re-absorption (Matthay, Folkesson & Clerici, 2002), causing what is termed clinically ARDS (Matthay, Ware & Zimmerman, 2012). Epithelial sodium channels (ENaC) located at the apical membrane regulate alveolar fluid reabsorption across the tight alveolar epithelium (Ji, Zhao, Chen, Shetty, Idell & Matalon, 2012). Reduced ENaC expression and activity are described in both injured human lungs ex vivo and in the animal models of lung injury (Matthay, Folkesson & Clerici, 2002). Decreased alveolar fluid clearance is confirmed in mice with deficient scnn1 genes. Plasminogen activators may decrease the severity of lung injury and pleural effusion (Karandashova et al., 2013) and may reduce the risk of death from ARDS and multiorgan failure from sepsis (Hardaway, Harke, Tyroch, Williams, Vazquez & Krause, 2001; Lim & Chin, 1999). Recently, our meta-analysis reports that fibrinolytic therapy might reduce lung injury, facilitate oedema fluid resolution, and reduced histologic lung injury score in animal models of acute lung injury (Liu et al., 2018). However, whether exogenously administered plasmin enhances the removal of oedema fluid via ENaC and related mechanisms, is not clear. We recently identified human ENaC as a novel target of urokinase (Ji, Zhao, Komissarov, Chang, Liu & Matthay, 2015). Based on the lesser specificity of plasmin substrates in amino acid sequence, it is a challenge to identify all of cleavage sites for plasmin, in particular for *in vivo* lifetime exposure to plasmin. On the other hand, whether or not plasmin could enhance reduced transepithelial fluid transport and restore impaired alveolar fluid clearance *in vivo* has not been studied. Therefore, we aimed to test the potential beneficial effects of plasmin delivered intratracheally on alveolar fluid clearance and determined proteolysis as post-translational mechanisms.

## 2. MATERIALS AND METHODS

### 2.1 Reagents

Human recombined plasmin (Molecular Innovation, Novi, Michigan, USA. Cat#: HPLM), two-chain urokinase plasminogen activator (Molecular Innovation, Novi, Michigan, USA. Cat#: UPA-HTC), dispase (Corning, USA.Cat.# 354235), DNase I (Sigma-Aldrich, St. Louis, Missouri, USA. Cat#: DN25), bovine serum albumin (Sigma-Aldrich, St. Louis, Missouri, USA. Cat#: A2153), biotin rat anti-mouse CD16/32 (BD, Pharmingen, USA. Clone 2.4G2. Cat#: 553143), biotin rat anti-mouse CD45 (BD, Pharmingen, USA. Clone 30-F11. Cat#: 553078), biotin rat anti-mouse Ter-119/erythroid TER-119 cells (BD, Pharmingen, USA. Cat# 553672), Dynabeads(tm) MyOne(tm) streptavidin T1 magnetic beads (Invitrogen, USA. Cat#: 65601), mouse IgG (Sigma-Aldrich, St. Louis, Missouri, USA. Cat# I5381), mouse laminin1 (Trevigen, USA. Cat#: 3401-010-02), tricaine-S (MS-222) (Western Chemical, Inc. USA), EZ-Link Sulfo-NHS-SS-Biotin (Thermo Scientific, USA. Cat#: 21331), anti-V5 mouse monoclonal IgG2a antibody (Invitrogen, USA. Cat#: R960-25), anti-HA high affinity rat monoclonal antibody (Roche USA. Clone 3F10. Cat#: 11867423001).

### 2.2 Animals and HCl instillation

Healthy, 8-12-week-old, male or female C57 BL/6 mice (cat#: 000664, Jackson Lab) were used. Animals were kept under pathogen-free conditions, and all experimental procedures were approved by the Institutional Animal Care and Use Committee of the University of Texas Health Science Center at Tyler. To create an *in vivo* model of lung injury, anesthetized mice were administered intratracheally (i.t.) 0.1M HCl in 2.5 μl·g^-1^ body weight followed by 30 μl·g^-1^ body weight of air (Nagase et al., 2000; Patel, Wilson & Takata, 2012). Control mice were administered the same volume of 0.9% NaCl and air.

### 2.3 *In vivo* alveolar fluid clearance (AFC) in mice

*In vivo* AFC rate was measured *in vivo* as previously described (Han et al., 2010; Patel, Wilson & Takata, 2012). Briefly, mice 4 hr post HCl instillation were anesthesized by ketamine (1.7 mg·ml^-1^) and xylazine (20 mg·ml^-1^), at 5ml/kg body weight by intraperitoneal injection. Then the mice were placed on a continuous positive airway pressure system delivering 100% O_2_ at 8 cmH_2_O. All animals were maintained at a temperature of 37 °C with a heating pad and ultrared bulb. An isosmotic instillate containing 5% bovine serum albumin was prepared with 0.9% NaCl as base solution. Control animals were intratracheally delivered with 50 μl the base solution. Plasmin (one group with 60 μg·ml^-1^ and another group with 100 μg·ml^-1^), a mixture of plasmin and equal amount of α2-antiplasmin (Pl + AP), and plasmin (60 μg·ml^-1^) in the presence of amiloride (1 mM) were intratracheally delivered. To maximize the collection of instilled BSA solution, the diaphragm was dissected post 30 min ventilation. Methodological concerns were addressed by using amiloride to inhibit water movement to confirm that measurements accurately reflected AFC. With the assumption that the concentration of instilled BSA proteins is not altered significantly by the presence of excess alveolar fluid and/or protein caused by HCl-induced damage. HCl may impair the blood-gas barrier function and causes a significant leakage of murine serum proteins into the air space. The endogenous BSA may lead to an overestimation of AFC rates. This potential bias was corrected by measuring murine albumin (cat#: EMA3201-1, AssayPro, St. Charles, MO) and bovine albumin (cat#: E10-113, Bethyl Laboratories, Montgomery, TX) separately using specific ELISAs following the manufacturer’s instructions (Wolk et al., 2008). Final AFC values were corrected by removing murine albumin present in the aspirate. In addition, we used an aliquot of the instillate that had been delivered into the lungs and removed within 5 min, rather than freshly prepared naive instillate *per se*. Any pre-existing oedema or leaking murine plasma proteins would be excluded for it dilutes both the control at time point 0 min (*P*_*i*_) and final samples (*P*_*f*_). The calculation of AFC was same as described for human lungs below.

### 2.3 *Ex vivo* human lung lobe studies

Discarded human lung lobes more than 5 cm away from tumor were collected 24 hr post-surgery with an approved exempt from the IRB committee of the China Medical University. The left and right lobes were chosen randomly upon availability. Lung lobes were ventilated with 100% O_2_ via a volume-controlled ventilator (model 683, Harvard Apparatus) and were passively rewarmed at 37 °C for 1 hr without perfusion in an incubator prior to measuring AFC. Plasmin (60 μg·ml^-1^) was added to the instillate (20 ml) and intrabronchially delivered. The instilled alveolar fluid was aspirated by applying gentle suction to the tracheal catheter with a 1-ml syringe. The BSA content of the alveolar fluid was measured with a 96-well microplate reader. AFC was calculated as follows: *AFC = (V*_*i*_ *- V*_*f*_*)/V*_*i*_ × *100*, where *V*_*i*_ and *V*_*f*_ denote the volume of the instilled and recovered alveolar fluid, respectively. *V*_*f*_ was obtained as *V*_*f*_ *= (V*_*i*_ × *P*_*i*_*)/P*_*f*_, where *P*_*i*_ and *P*_*f*_ represent protein concentration of instilled and collected fluid.

### 2.4 Primary type 2 alveolar epithelial cell (AT2) culture and bioelectrical measurements

Mouse AT2 cells were isolated from both wt and uPA knockout (*plau*^*-/-*^) strains of C57BL/6 mice (cat#: 000664 and 002509, respectively. Jackson Laboratory, USA) with a modified protocol as described (Demaio et al., 2009; Zhao et al., 2019). Briefly, lungs of euthanized mice were removed and incubated in dispase for 45 min, and then were gently teased and incubated in DMEM/F-12 + 0.01% DNase I for 10 min. Cells were passed through a serial of Nitex filters (100, 40, 30, and 10 microns; Corning, USA) and centrifuged at 300 g for 10 min. Resuspended cells were biotinylated with CD16/32 (0.65 μg·10^−6^cells), CD45 (1.5 μg·10^−6^cells) and Ter/119 (10 μl)) antibodies and then incubated with streptavidin-coated magnetic particles (2.5 μg·10^−6^cells). Fibroblasts were removed by a 2 hr incubation in a mouse IgG coated plastic culture dish. Upon passing the request for both viability (>90%) and purity assays (>95%), mouse laminin 1 precoated transwells (cat#: 3413, Corning Costar, USA) were seeded with AT2 cells (10^6^ cells per cm^2^). Culture medium was replaced post 72 hr and then every 48 hr. Transepithelial resistance (R_TE_) and potential difference (V_TE_) were measured using an epithelial voltohmmeter (World Precision Instrument, USA). Equivalent short-circuit current (I_EQ_) values were calculated as the ratio of V_TE_ / R_TE_.

### 2.5 Construction of ENaC mutants

Deletion and site-directed mutants were generated in human ENaC cDNAs cloned into a pGEM HE vector using the QuikChange II Site-Directed Mutagenesis kit (Stratagene) (Ji & Benos, 2004; Molina et al., 2011). cRNAs of human α, β, and γ ENaC subunits were prepared as described previously (Ji, Parker, Langloh, Fuller & Benos, 2001). HA and V5 tags were introduced to the N and C-termini of α and γ ENaC subunits, respectively.

### 2.6 Oocyte expression and voltage clamp studies

*Xenopus* oocytes were surgically removed from appropriately anesthetized adult female *Xenopus laevis* (cat#: HCG IMP FM, hCG injected Mature Female *Xenopus* laevis, Xenopus Express) as described (Zhao et al., 2019). Briefly, the ovarian tissue was removed from frogs under anaesthesia by 0.1% of Tricaine through a small incision in the lower abdomen. Ovarian lobes were removed and digested in OR-2 calcium-free medium (in mM: 82.5 NaCl, 2.5 KCl, 1.0 MgCl_2_, 1.0 Na_2_HPO_4_, and 10.0 HEPES, pH 7.5) with the addition of 2 mg·ml^-1^ collagenase (Roche, Indianapolis). Defolliculated oocytes were injected with ENaC cRNAs into the cytosol (25 ng per oocyte in 50 nl of RNase-free water) and incubated in regular OR-2 medium at 18 °C. The two-electrode voltage clamp technique was used to record whole-cell currents 48 hr post injection. Oocytes were impaled with two electrodes filled with 3M KCl, having resistance of 0.5 - 2 MΩ. A TEV-200A voltage clamp amplifier (Dagan) was used to clamp oocytes with concomitant recording of currents. Two reference electrodes were connected to the bath. The continuously perfused bathing solution was ND-96 medium (in mM: 96.0 NaCl, 1.0 MgCl_2_, 1.8 CaCl_2_, 2.5 KCl, and 5.0 HEPES, pH 7.5). Whole-cell currents were recorded as previously reported (Ji et al., 2006). Experiments were controlled by pCLAMP 10.7 software (Molecular Devices), and currents were continuously monitored at an interval of 10 sec. To analyse amiloride inhibition, we waited until the current level reduced and reach a plateau and last for at least 1 min. after amiloride application to the bath Data were sampled at the rate of 200 Hz and filtered at 50 Hz.

### 2.7 Biotinylation and immunoblotting

Biotinylation experiments were adapted from previous publications (Haerteis, Krappitz, Diakov, Krappitz, Rauh & Korbmacher, 2012; Haerteis, Krueger, Korbmacher & Rauh, 2009), using 20-40 oocytes per group. In some experiments, oocytes were preincubated either in ND-96 solution or low sodium solution (1 mM NaCl, 96 mM NMDG). Oocytes were incubated in freshly prepared biotinylation buffer (1.5 mg·ml^-1^ EZ-Link Sulfo-NHS-SS-Biotin in DPBS solution from Hyclone, pH 8.0) for 30 min at room temperature with gentle agitation. The biotinylation reaction was stopped by washing the oocytes three times for 5 min each with quenching buffer (in mM: 192 mM glycine and 25 mM Tris-Cl, pH 7.5). Subsequently, the oocytes were incubated in ND-96 solution or supplemented with 10 μg·ml^-1^ human plasmin for 60 min or designated periods for time-dependent study. After washing the oocytes three times with ND-96 solution, treated cells were lysed by passing them through a 27-gauge needle in lysis buffer (in mM: 500 mM NaCl, 5 mM EDTA, 50 mM Tris, 1% Triton X-100, 1% Igepal CA-630, pH 7.4) and supplemented with Complete Mini EDTA-free protease inhibitor cocktail (Roche, 04693159001) according to the manufacturer’s instructions. The lysates were incubated in a shaker for 1 hr and centrifuged at 16,000 ×g for 15 min at 4 °C. Supernatants were transferred to 1.5-ml tubes (Eppendorf). Biotinylated proteins were precipitated with 50 μl of pre-washed high capacity neutravidin agarose resin (Pierce, 29204). After overnight incubation at 4 °C with overhead rotation, supernatants were removed, and beads were washed three times with lysis buffer containing protease inhibitors. 50 μl of 2× SDS-PAGE sample buffer (Pierce, 39001) was added to the beads. Samples were boiled for 5 min at 95°C, centrifuged for 1 min and loaded on a 7.5% SDS-PAGE gel. To detect small peptides by anti-HA antibody, samples were run on a 16.5% Tris-Tricine gel (BioRad, 4563063) for some experiments. To detect γ ENaC fragments, the membrane blots were blocked in 5% blocking buffer (5% non-fat dry milk, Bio-Rad, in TBST) for 1 h at room temperature. Then, anti-V5 or anti-HA monoclonal) antibodies were added to the samples (1:5,000 and 1:1,000 dilution, respectively). Our and other groups have demonstrated that non-ENaC bands could be detected in parental oocytes or oocytes expressing ENaC constructs without these artificial flags (Carattino et al., 2014; Ji, Zhao, Komissarov, Chang, Liu & Matthay, 2015; Passero, Mueller, Myerburg, Carattino, Hughey & Kleyman, 2012; Passero, Mueller, Rondon-Berrios, Tofovic, Hughey & Kleyman, 2008). Horseradish peroxidase-labelled secondary antibodies (Jackson Immunoresearch) were used (1:10,000). Chemiluminescence signals were detected using ECL Plus (Millipore).

### 2.8 *In silico* prediction of plasmin cleaved sites in α, β, γ, and δ ENaC subunits

Specific cleavage sites for human plasmin confirmed with phage substrate display and solution phase fluorogenic peptide microarray (Gosalia, Salisbury, Maly, Ellman & Diamond, 2005; Hervio, Coombs, Bergstrom, Trivedi, Corey & Madison, 2000) were applied. Another substrate motif from P4 to P4’ was from the getMerops of the SitePrediction server (Verspurten, Gevaert, Declercq & Vandenabeele, 2009). Default settings of the server were used. The predicted sites must meet these criteria: 1) the cleavage sites are located at the ectodomain of ENaC, 2) the size of predicted C-terminal fragments is similar to that on Western blots or smaller considering potential multiple proteolysis, 3) the P1 protein must be either Arg (R) or Lys (K), 4) average score is >1, 5) specificity is > 95%, and 6) predicted by applying both optimized substrate sequences.

### 2.9 Mass spectrometry analysis for plasmin cleaved fragments

Three pure peptides (with a purity >95%) with a continuous sequence from amino acid residue T121 to A190 of the human γ ENaC proteins were synthesized by the GenScript (NJ, USA). The N- and C-termini were acetyled and amidated, respectively. The sequence of each peptide was show in Figure 8a. The synthesized peptides were dissolved in 400 μl DPBS (1.0 mg·ml^-1^), which was digested by adding 10 μg·ml^-1^ plasmin for 30 min at room temperature. Trifluoroacetic acid (0.1%) was used to inactivate cleavage process. The samples were analysed by the University of Texas Southwest Medical Center Proteomics Core. Data were then analysed with the Proteoe Discoverer 2.2 software (Thermo Fisher, USA) based on a tryptic digestion with a maximum of 6 missed cleavages at amino acid residues R and K. F and V sites are searched by the strategy for non-specific cleavage. The frequency of the fragments with cleaved end of a specific amino acid residue is the sum of peptide spectral matches for both termini.

### 2.10 Statistical analysis

All results were presented as mean ± S.E. ENaC activity is the difference of the total and amiloride-resistant current fractions. One-way ANOVA computation combined with the Bonferroni test was used to analyse the difference of the means for significance. P < 0.05 was considered significant.

### 2.11 Nomenclature of targets and ligands

Key protein targets and ligands in this article are hyperlinked to corresponding entries in http://www.guidetopharmacology.org, the common portal for data from the IUPHAR/BPS Guide to PHARMACOLOGY (Harding et al., 2018), and are permanently archived in the Concise Guide to PHARMACOLOGY 2017/18 (Alexander et al., 2017).

## 3 RESULTS

### 3.1 Plasmin increases alveolar fluid clearance in healthy human lungs

The beneficial effects of plasminogen activators on septic respiratory dysfunction imply that plasmin might improve AFC via the ENaC pathway (Chen et al., 2014; Liu et al., 2018). To address this issue, the effects of plasmin on ENaC-mediated fluid resolution were measured in ventilated *ex vivo* human lung lobes (Figure 1a). Plasmin significantly increased overall AFC value from 14% in control lobes to 37% in treated lobes (P < 0.05) in 1 hr. Approximately 75% of the AFC rate was ENaC-dependent, the fraction that was inhibited by amiloride. As reflected by the proportion of amiloride-inhibitable AFC, plasmin stimulated ENaC activity up to 3 times (Figure 1b).

**FIGURE 1.**
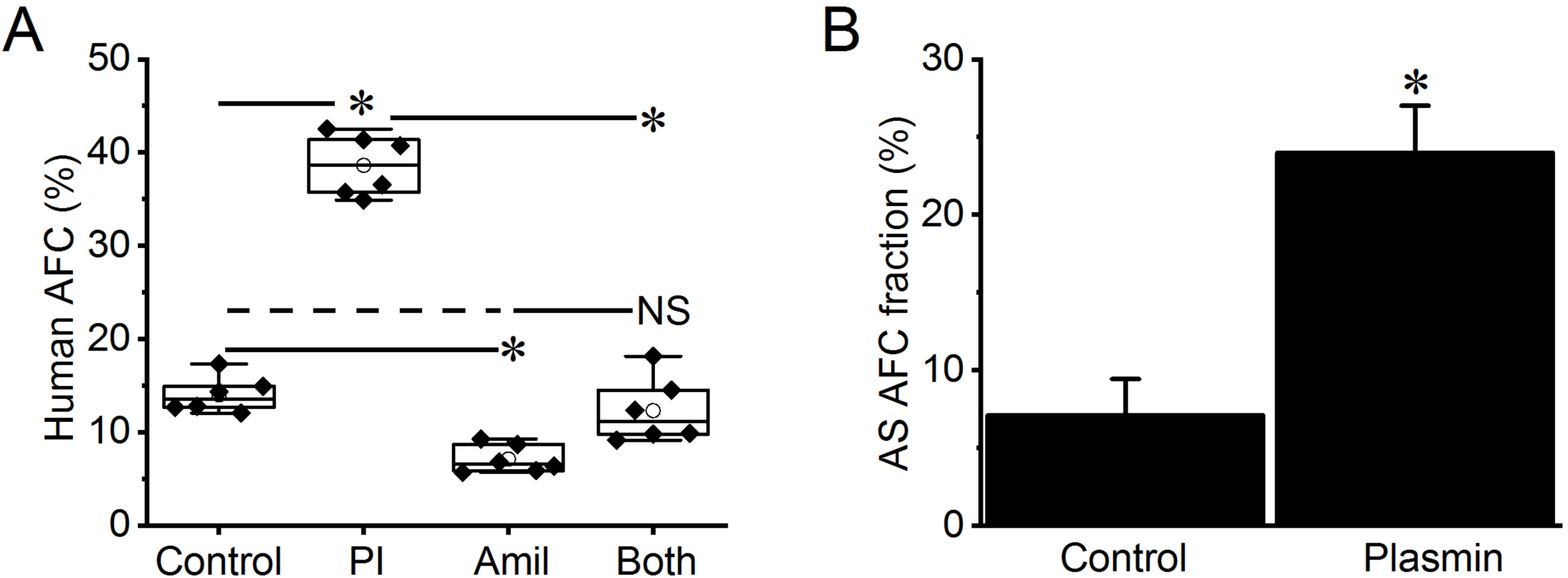
Intratracheal administered plasmin up-regulates alveolar fluid clearance in normal human lungs. *a*. Total *ex vivo* 60-min alveolar fluid clearance (AFC_60_) in the presence and absence (Control) of plasmin (Pl, 60 μg·ml^-1^, 0.7 μM·L^-1^) and/or amiloride (Amil, 1 mM·L^-1^). Adjacent human lung lobes within one biopsy specimen were selected and AFC values were measured in parallel for both control and treated groups. Data were presented as median (filled circle), mean (horizontal line), 25%, 75% percentile, and standard error. * P < 0.05 vs those in the absence of plasmin. NS, not significant vs both Control and Amil. n = 24. *b*. Amiloride sensitive (AS) fraction of AFC was computed as ENaC contribution in human lung lobes instilled with PBS only (Control) and plasmin. * P < 0.05. n = 24.

### 3.2 Plasmin increases alveolar oedema fluid clearance in the mouse model of injured lungs

We postulated that plasmin could restore the impaired ENaC function in a mouse model of gastric acid aspiration (Nagase et al., 2000). Intratracheal administration of two boluses of plasmin 1 hr post HCl instillation significantly reduced lung water content of injured mouse lungs (Figure 2a). In contrast, deactivation of the catalytic activity of plasmin with specific α2-antiplasmin or amiloride did not alter the wet/dry ratio significantly. Correspondingly, *in vivo* AFC rate was restored close to a near normal level by proteolytically active plasmin but not by the mixture of plasmin with either α2-antiplasmin or amiloride (Figure 2b). Together with the results in *ex vivo* human lungs, these data suggest that plasmin may resolve oedema fluid via the ENaC pathway in injured lungs.

**FIGURE 2.**
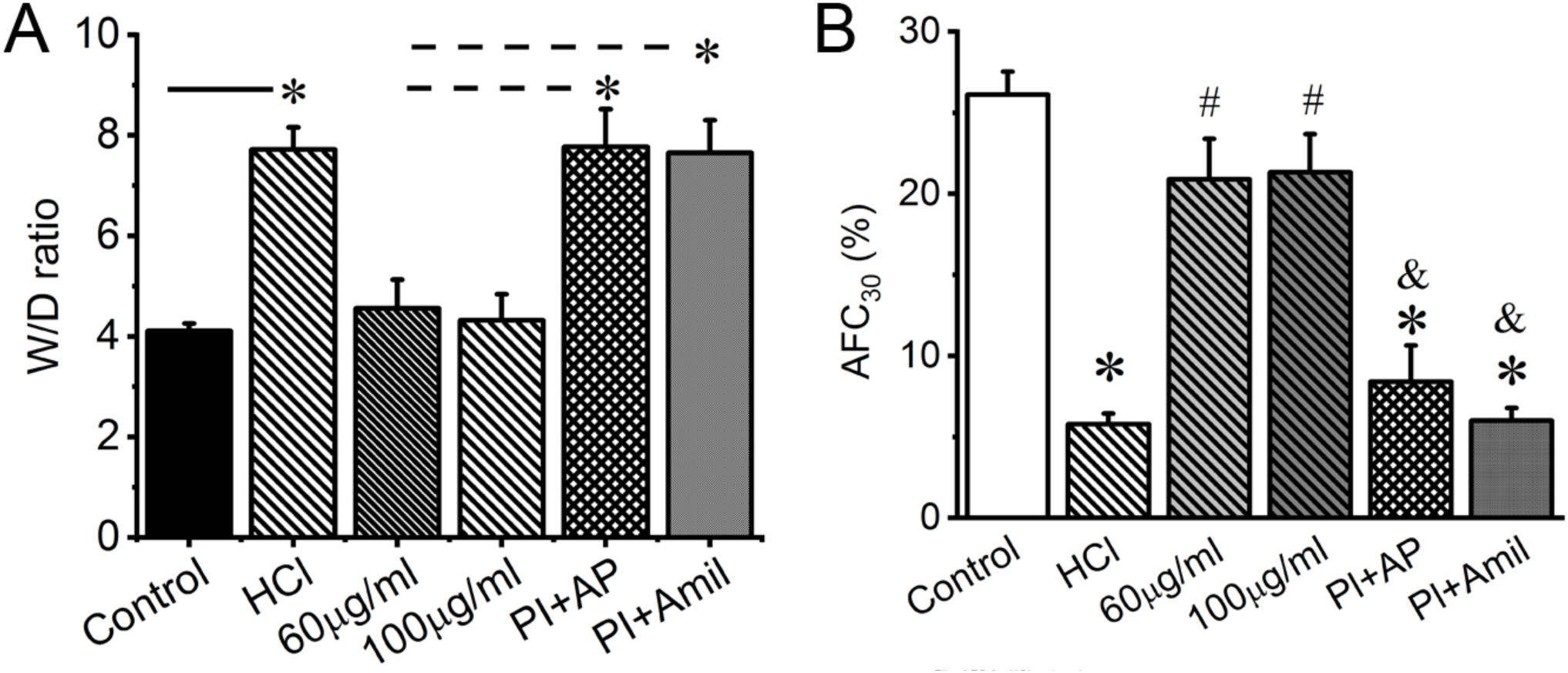
Plasmin stimulates alveolar fluid clearance in acid aspiration injured mouse lungs. *a*. Lung water content (wet/dry ratio). Mice were intratracheally instilled 1N HCl (50 μl) as a model of gastric acid aspiration induced lung injury. Control animals were intratracheally delivered with 50 μl PBS. Plasmin (60 μg·ml^-1^ and 100 μg·ml^-1^) and a mixture of plasmin and equal amount of α2-antiplasmin (Pl + AP) were intratracheally delivered 4 hr post HCl instillation. Lungs were dissected and subjected to weigh before and after dehydration. *P < 0.05. Dashed line indicates the difference vs more than one groups below. *b*. Plasmin restores *in vivo* 30-min (AFC_30_) in mice injured by HCl. n = 20. * P < 0.05 vs Control. # P < 0.05 vs HCl group. & P < 0.05 vs plasmin only groups.

### 3.3 Plasmin stimulates deficient ENaC activity in primary AT2 monolayers

Plasmin exhibited multifaceted effects on membrane-arched receptors and enzymatic substrates in epithelial and nonepithelial cells (Castellino & Ploplis, 2005; Schaller & Gerber, 2011). To further confirm the stimulatory effects of plasmin on ENaC function located at the apical membrane of alveolar epithelial cells, we applied plasmin to polarized primary AT2 cell monolayers. As shown in Figure 3a-b, a significant suppression in ENaC activity accompanied by an increment in transepithelial resistance was seen in *plau*^*-/-*^ deficient monolayers, an *in vitro* model of lung injury mimicking eliminated fibrinolytic activity in injured lungs. The reduced ENaC function was recovered by a bolus of plasmin (Figure 3c). Combined with our previous studies in *plau*^*-/-*^ mice and other groups’ observations on the regulation of ENaC by plasmin (Chen et al., 2014; Haerteis, Krappitz, Diakov, Krappitz, Rauh & Korbmacher, 2012; Ji, Zhao, Komissarov, Chang, Liu & Matthay, 2015; Passero, Mueller, Rondon-Berrios, Tofovic, Hughey & Kleyman, 2008), we set up to examine the post-translational mechanisms for plasmin to augment human αβγENaC channels heterologously expressed in *Xenopus* oocytes.

**FIGURE 3.**
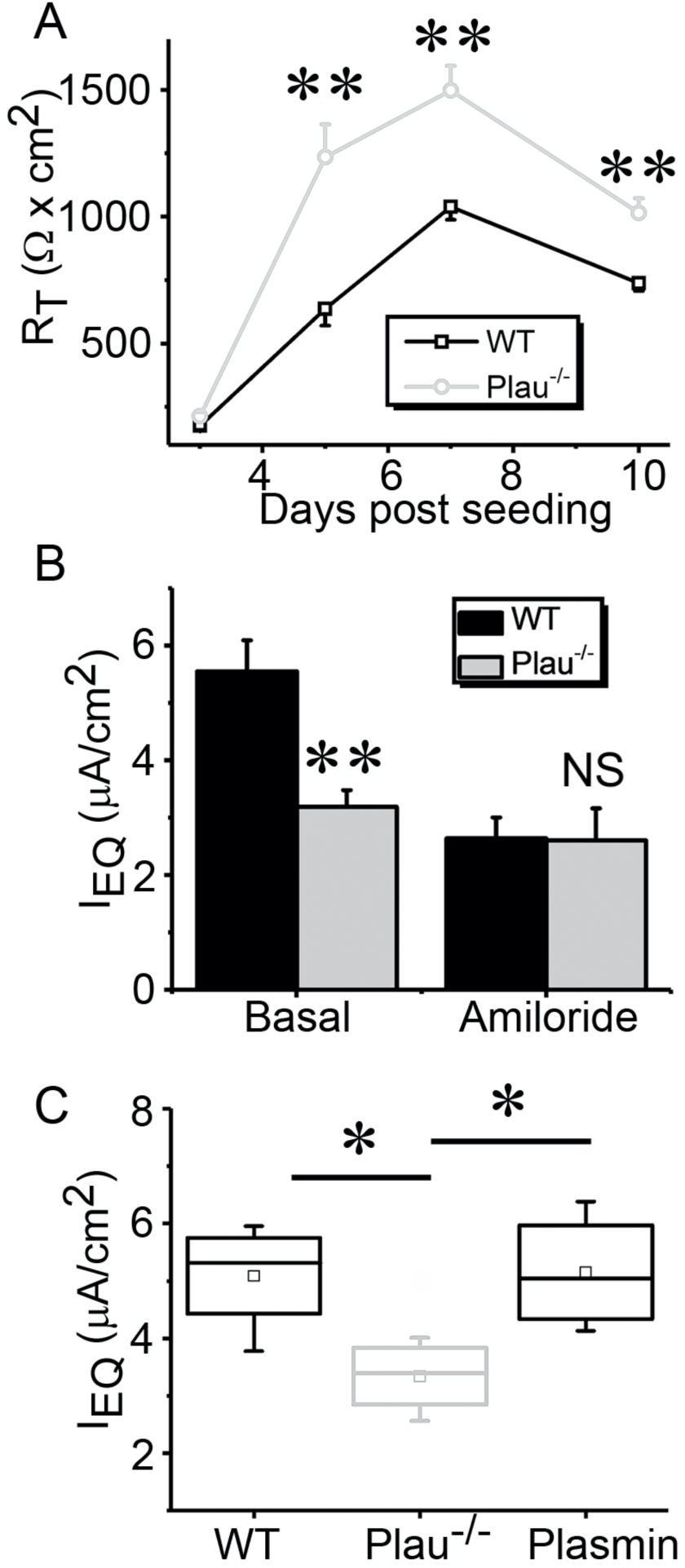
Plasmin restores deficient ENaC activity in uPA deficient alveolar type 2 (AT2) epithelial monolayer cells. *a*. Transepithelial resistance in wt and *plau*^*-/-*^ monolayers up to 10 days post seeding. ** P < 0.01 vs wt cells. n = 6. *b*. Equivalent short-circuit (IEQ) in polarized AT2 monolayers cultured at the air-liquid mode. IEQ = ET/RT. ET is the transepithelial potential difference, and RT is the resistance across the monolayer read by an EVOM meter. Data were presented as mean value and standard error. NS, not significant, ** P < 0.01 vs wt group. n = 6. *c*. Effects of plasmin on IEQ in polarized AT2 monolayers. Data were presented as median (filled circle), mean (horizontal line), 25%, 75% percentile, and standard error for wild type (wt), *plau* knockout (*plau*^*-/-*^), and addition of plasmin 20 μg·ml^-1^. * P < 0.05. n = 5-7.

### 3.4 γENaC is targeted by plasmin

Plasmin is generated from plasminogen, a substrate of both tissue-type and urokinase-like plasminogen activators (tPA and uPA, respectively). To examine the effects of plasmin on ENaC activity, we incubated cells heterologously expressing human αβγENaC with plasmin for 1 hr with uPA as a positive control (Ji, Zhao, Komissarov, Chang, Liu & Matthay, 2015) (Figure 4a). Plasmin activated ENaC currents to approximate 35 μA from 5 μA, which was more potent than tc-uPA. In sharp contrast, the mixture of plasmin and α2-antiplasmin did not affect ENaC current. Plasmin activated human heterologous αβγENaC activity in a time-dependent manner. The channel activity was activated maximally in 1 hr (Figure 4b). Subsequently, ENaC function declined but still maintained at a level significantly greater than controls within 8 hr. To identify what subunit was targeted by plasmin, the stimulatory effects of plasmin on ENaC channels comprised of α subunit alone, α + β subunits, and α+ γ subunits were analysed with αβγ channels as a positive control (Figure 4c). Both current amplitude and fold of αγENaC increased by plasmin significantly, implying that γ subunit could be targeted by plasmin (Figure 4d). In contrast, the currents of α alone and α + β ENaC channels were not significantly altered by plasmin. Further, with chymotrypsin as a positive control, we examined potential proteolytic cleavage of α and γ ENaC subunits with immunoblotting assays (Figure 4e-h). Both N- and C-terminal fragments were detected by anti-HA and anti-V5 monoclonal antibodies, respectively. As reported previously (Carattino, Sheng, Bruns, Pilewski, Hughey & Kleyman, 2006), full-length αENaC subunit (90 kDa) was cleaved by endogenous proteases (i.e., furin) into two fragments in control group (Figure 4e-f). Neither plasmin nor chymotrypsin altered furin-cleaved V5- (64 kDa,) and HA-tagged fragments (26 kDa). In sharp contrast, plasmin and chymotrypsin reduced the furin-cleaved C-terminal fragment (80 kDa) of full-length γENaC proteins (90 kDa) to 70 kDa (Figure 4g). The N-terminal fragment (26 kDa) as recognized by anti-HA antibody was not affected by both plasmin and chymotrypsin (Figure 4h). Consistent with our previous studies, serine proteases are unable to cleave full-length ENaC (Ji, Zhao, Komissarov, Chang, Liu & Matthay, 2015). These functional and immunoblotting data indicate that plasmin may specifically activate human αβγ channels by targeting γ subunits proteolytically, and that plasmin as well as chymotrypsin may not be able to cleave the HA-tagged N-terminal fragments of both α and γ subunits cut off by furin.

**FIGURE 4.**
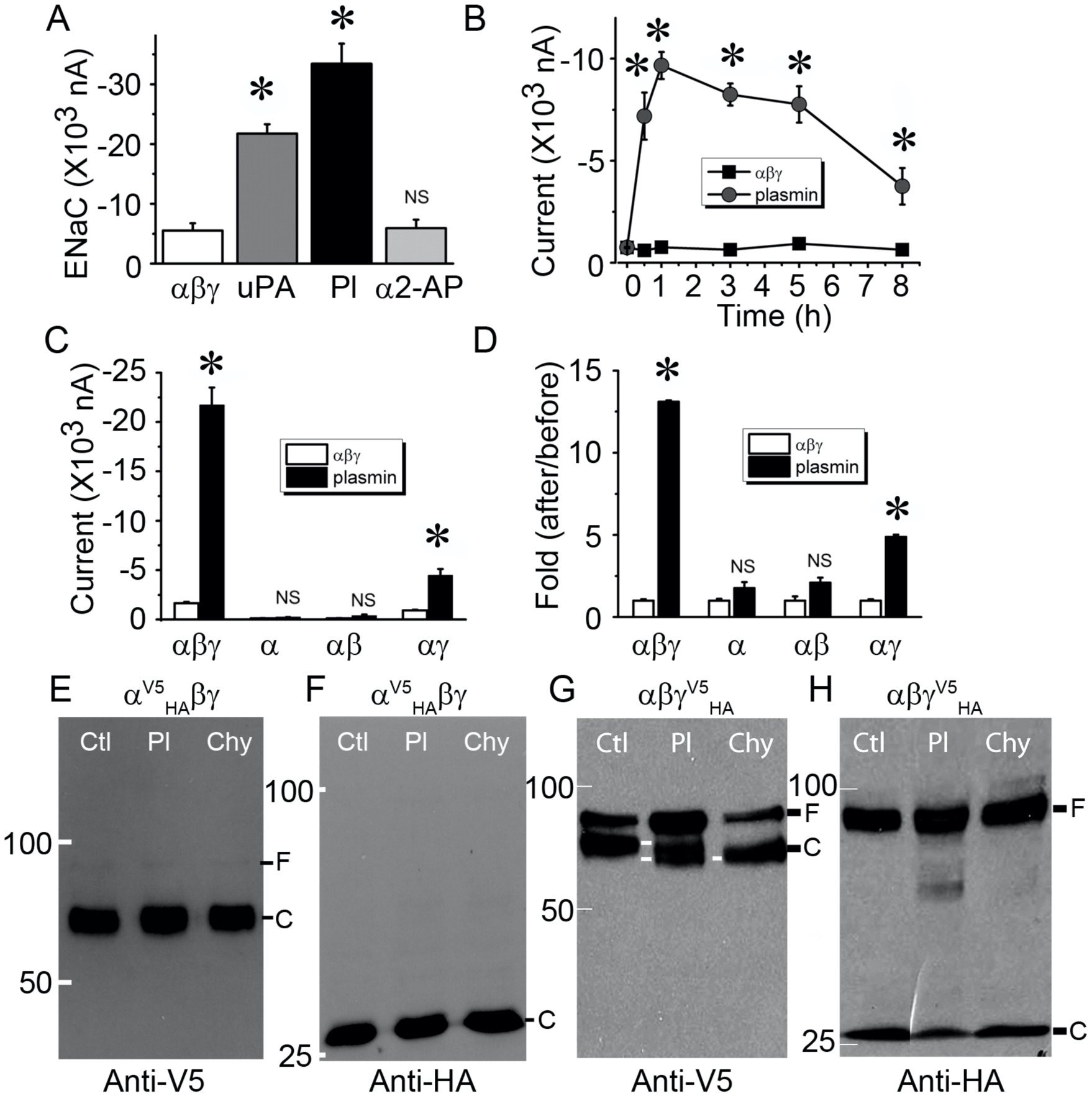
Plasmin specifically activates human αβγ ENaC channels expressed in *Xenopus* oocytes. *a*. Amiloride-sensitive sodium currents (ENaC) were digitally sampled at the membrane potential of −100 mV in cells heterologously expressing human αβγ ENaC, incubated with two-chain urokinase plasminogen activator (uPA, 10 μg·ml^-1^, 30 min at room temperature), plasmin (Pl, 10 μg·ml^-1^), and a complex of plasmin (Pl, 10 μg·ml^-1^) and α2-antiplasmin (α2-AP, 10 μg·ml^-1^). n = 20 per group. NS, not significant, * P < 0.05 vs controls in the absence of fibrinolytic molecules. *b*. Time course of ENaC activation by plasmin. ENaC activity in the set of oocytes was recorded at designated time points post exposure to plasmin (closed circle). n = 6 per group. * P < 0.05 vs the corresponding time point of controls (closed square). *c*. Effects of plasmin on the current levels of α, α + β (αβ), and α + γ (αγ) channels. n = 10-12. NS, not significant, * P < 0.05 vs the corresponding controls (open bar) in the absence of plasmin. *d*. Normalized incremental fold of ENaC activity. Ratio of current amplitudes recorded after over before addition of plasmin or perfusate was computed as fold of increase in channel activity. NS, not significant, * P < 0.05 vs controls. *e & f*. Detection of full-length (F) and cleavage (C) of α ENaC by plasmin. HA (attached to the N-terminal tail) and V5 (attached to the C-terminal tail) tagged α subunit (α^V5^HA) was co-expressed with βγ subunits complementarily. Biotinylated plasma membrane proteins were run on a 7.5% SDS-PAGE gel and probed with anti-V5 (*e*) and anti-HA (*f*) monoclonal antibodies. Lanes from left to right were loaded with plasma membrane proteins of cells without plasmin treatment (Ctl), pretreated with plasmin (Pl, 10 μg·ml^-1^ for 30 min at room temperature), and chymotrypsin (Chy, 10 μg·ml^-1^) as a positive control. *g & h*. Cleavage of HA and V5 tagged γ ENaC (γ^V5^HA) by plasmin as recognized by anti-V5 (*g*) and anti-HA (*h*) antibodies. Cleaved bands are labeled with white lines at the same level. These experiments were repeated at least four times with similar observations.

### 3.5 Plasmin-mediated regulation of deletion and site-directed mutants missing putative cleavage domains

Three consensus domains at the extracellular loop for proteases to cleave ENaC proteins have been proposed (Kleyman, Carattino & Hughey, 2009; Planes & Caughey, 2007; Rossier & Stutts, 2009). We attempted to find potential putative tracts responsible for plasmin-mediated cleavage in both α and γ subunits. As shown in Figure 5a, removal of the first (aa173-178) and second (aa201-204) putative cleavage domains in the finger of α ENaC slightly but not significantly reduced and augmented the stimulation of plasmin on ENaC function, respectively. By comparison, deletion of the third consensus cleavage tract (aa432-444) in the thumb markedly eliminated plasmin-induced up-regulation of the channel activity (Table 1 and Figure 5a). Similarly, all of three deletion mutants of γ ENaC subunit almost lost their responses to plasmin (Figure 5b). In sharp contrast, plasmin-cleaved C-terminal peptides with variant sizes were seen for these γ deletion mutants (Figure 5c-f). Intriguingly, this process might not be furin cleavage-dependent because plasmin was still able to cleave γΔ131-138, which lost furin cleavage sites-R135K136 (Figure 5d). Plasmin was reported to potentially cleave both human and mouse γENaC subunit (Haerteis, Krappitz, Diakov, Krappitz, Rauh & Korbmacher, 2012; Passero, Mueller, Rondon-Berrios, Tofovic, Hughey & Kleyman, 2008). Mutation of mouse γ K194 significantly suppressed acute activation of heterologous mouse αβγ ENaC channels in minutes (Passero, Mueller, Rondon-Berrios, Tofovic, Hughey & Kleyman, 2008). Substitution of the corresponding site in human γ ENaC subunit (γK189A) with alanine, however, could not prevent the stimulatory effects of plasmin on human αβγK189A channels (Figure 5g). Instead, both the domains for mouse plasmin- and prostasin-induced cleavage in human γ ENaC subunit termed γ5A (RKRK^178-181^AAAA + K189A) were crucial for plasmin to acutely activate human αβγ function (Haerteis, Krappitz, Diakov, Krappitz, Rauh & Korbmacher, 2012). Surprisingly, plasmin still increased the current amplitude of this αβγ5A channels approximate 9 and 20 times in 1 and 24 hr, respectively (Figure 5h). Meanwhile, the plasmin-cleaved C-terminal fragments were recognized by anti-V5 antibody (Figure 5i), suggesting additional new cleavage sites exist beyond these 5 amino acid residues for long-term exposed plasmin.

**Table 1.**
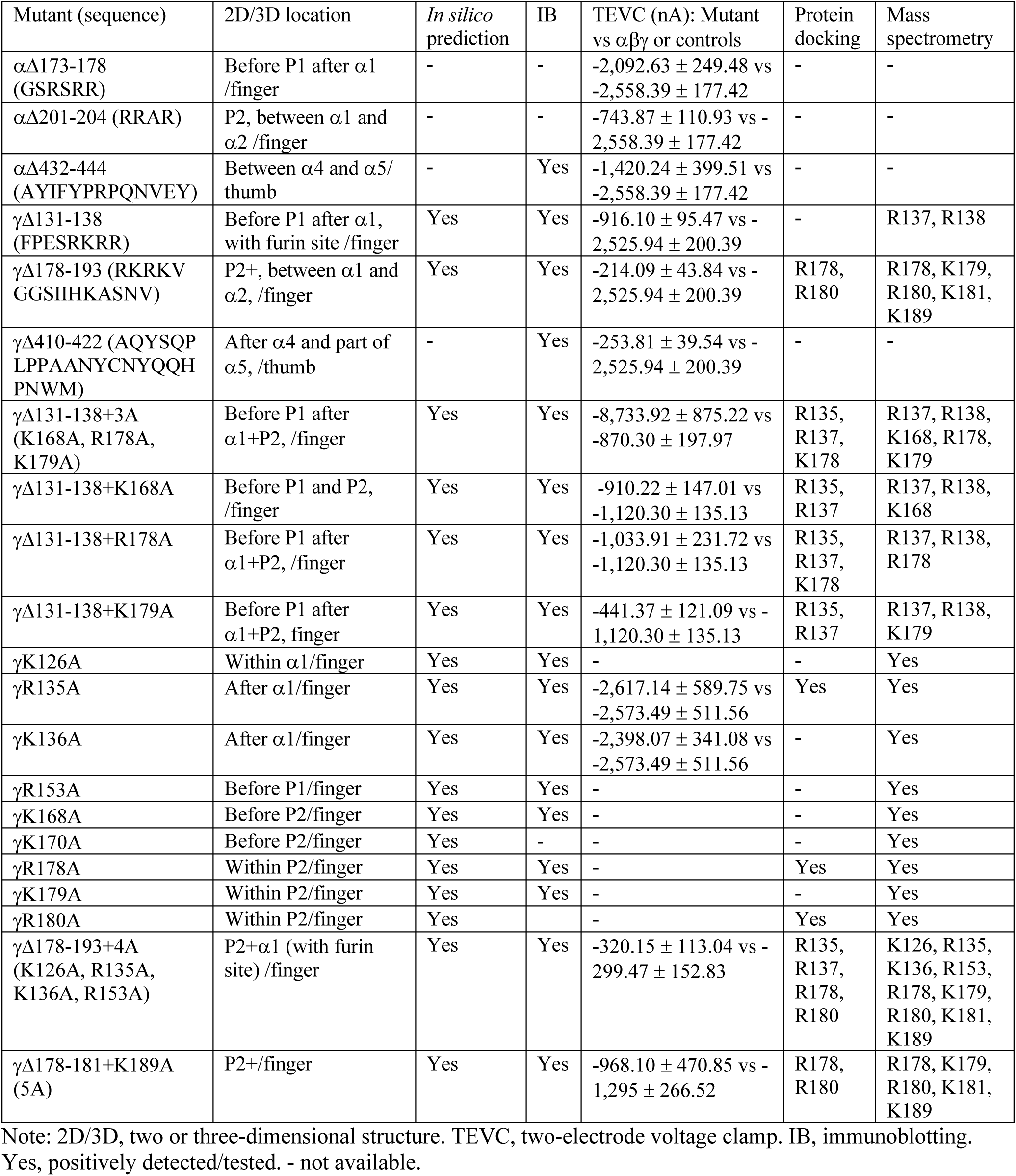
Characteristics of ENaC mutants. All of deleted and single or multiple directed mutants are included for predicted location in the 2D/3D cryo-EM model of truncated human ENaC (Noreng, Bharadwaj, Posert, Yoshioka & Baconguis, 2018) and experimental evidence.

**FIGURE 5.**
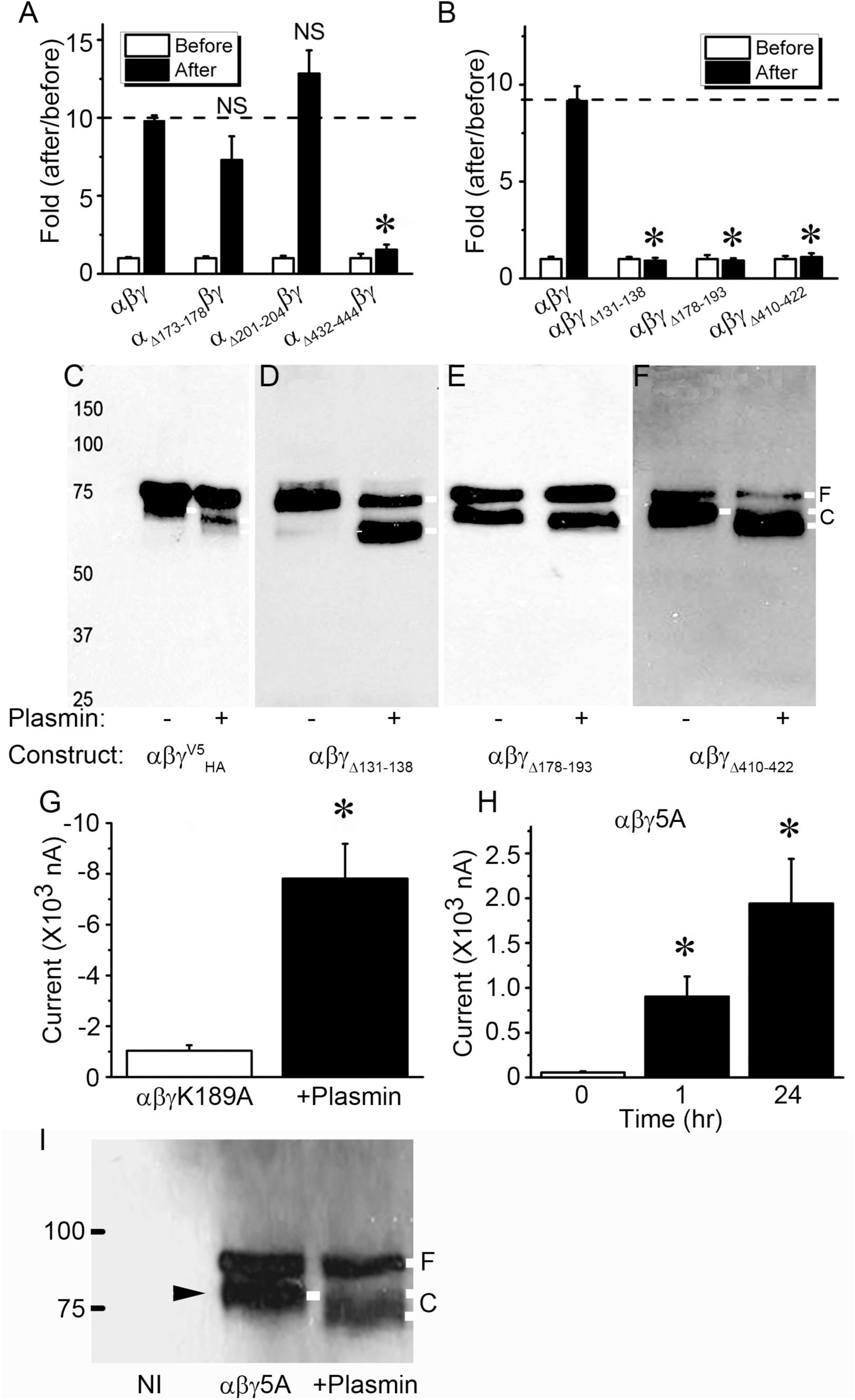
Effects of plasmin on α and γ ENaC mutants missing consensus cleavage domains. *a*. Basal ENaC currents were recorded in oocytes expressing αβγ ENaC and three deletion mutants lack of putative cleavage domains, namely, αΔ173-178βγ, αΔ201-204βγ, and αΔ432-444βγ. The activated currents were collected again 1 hr post incubation with plasmin (10 μg·ml^-1^, 30 min at room temperature) at the room temperature. The activated current levels were normalized to that basal currents. NS, not significant, *P < 0.05 vs αβγ ENaC. n = 19 - 26. *b*. Activation of three γ deletion mutants of αβγΔ131-138, αβγΔ178-193, and αβγΔ410-422 by plasmin. *c-f*. Immunoblotting detection of cleavage in full-length and three deletion γ mutants. The most left lane is protein marker. Untreated cells were controls (-). Endogenous furin cleaved band in the absence of plasmin and plasmin-cleaved band in the presence of plasmin (+) were labeled with white lines. These blots represent four experiments with similar observations. *g*. Plasmin activates the activity of mouse mutant (αβγK189A) for plasmin cleavage site. Whole-cell currents were digitized and then repeated 1 hr post incubation with plasmin (10 μg·ml^-1^). n = 8. * P < 0.05. *h*. Plasmin stimulates the activity of a human mutant (αβγ5A) for plasmin cleavage (K189 and 178RKRK181 were substituted with alanine). Measurements of whole-cell currents were performed 1 and 24 hr after exposure to plasmin. n = 22, * P < 0.05 vs those before addition of plasmin. *i*. Western blot assay of biotinylated plasma membrane proteins an anti-V5 monoclonal antibody. Plasma membrane proteins from noninjected oocytes (NI), cells expressing αβγ ENaC (Control), and cells pretreated with plasmin (Plasmin) were loaded on a 7.5% SDS-PAGE gel. Furin and plasmin-cleaved proteins are pointed out with black and white rightward arrowhead, respectively. Similar results were observed in four blots.

### 3.6 Identification of novel critical domains for activation of ENaC by plasmin

To identify substrate-like motifs in the γENaC proteins, we carried out *in silico* predictions with the SitePrediction server with two different strategies (Figure 6a). Because plasmin cleaved both full-length and furin-catalysed proteins as shown in Figure 5d, we simplified our study by using the furin-site deletion mutant, γΔ131-138 as a base construct to prepare a new triple mutant, namely, γΔ131-138 + 3A (K168A, R178A, K179A). These three predicted sites were top ranked by the SitePrediction server. As shown in Figure 6b, the activity of this mutant was not activated by plasmin in both 1 and 24 hr. The cleaved C-terminal fragments were not recognized compared with that of αβγΔ131-138 (Figure 6c). Moreover, three single point mutants were made: γΔ131-138 + K168A, γΔ131-138 + R178A, γΔ131-138 + K179A. The activity of these site-directed mutants was no longer elevated by plasmin (Figure 6d). Immunoblotting assays showed a significant reduction in the fraction of cleaved proteins accompanied by increased full-length bands (Figure 6e). Similarly, the densitometry ratio of cleaved bands over full-length bands was marked reduced for these single site-directed mutants (Figure 6f).

**FIGURE 6.**
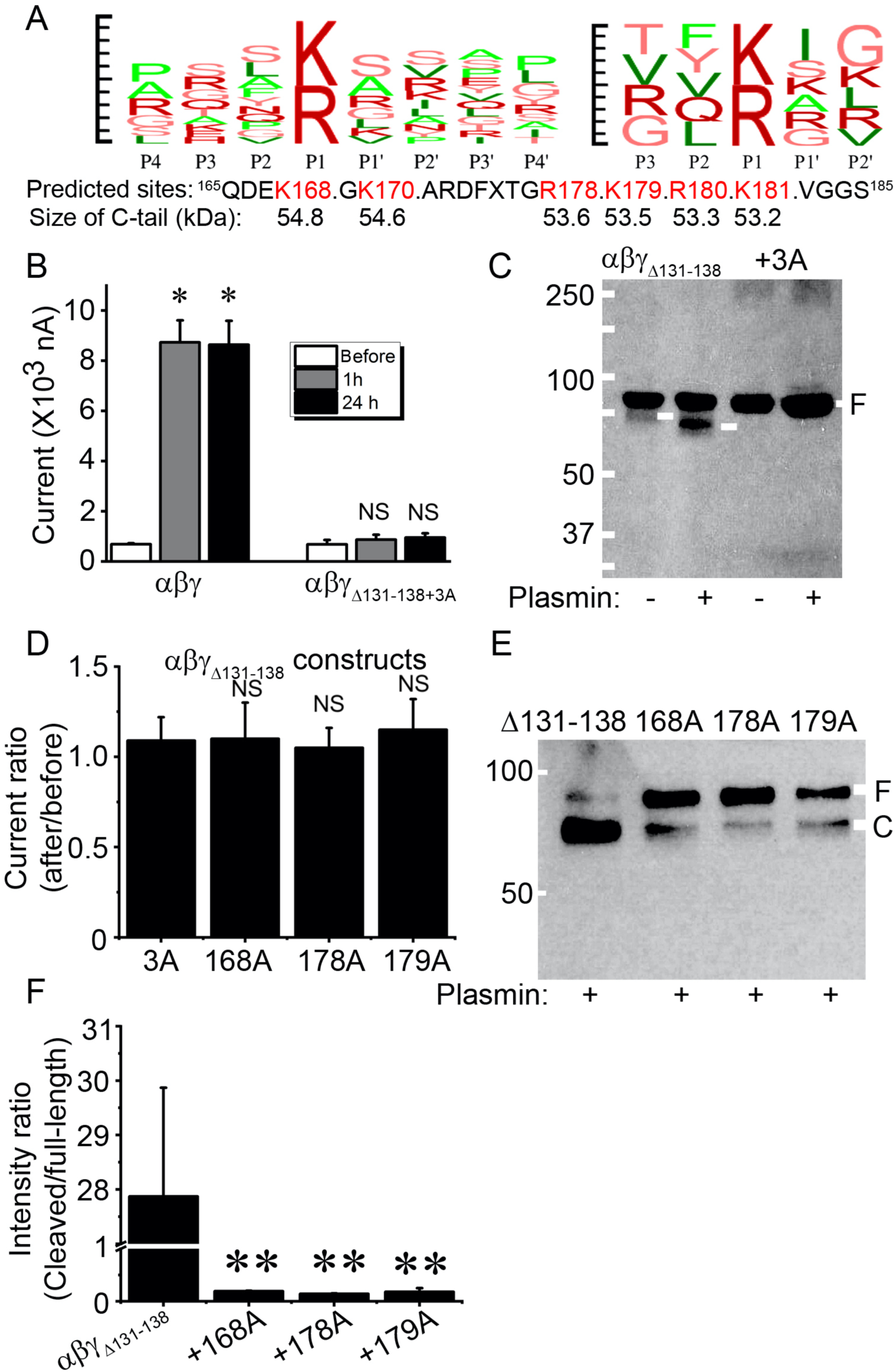
Potential proteolysis domains for plasmin beyond the putative cleaving motif (131-138) in γ ENaC subunit. *a. In silico* prediction with the database for plasmin-specific motifs. Left, a combination (logo) of reported 92 substrate motifs for plasmin from P4 to P4’. Right, a logo of input substrate sequences for plasmin from P3 to P2’. Top 10 predicted sites and the size of C-terminal fragments by both strategies are listed below. Default setup was used for running the SitePrediction server (Gosalia, Salisbury, Maly, Ellman & Diamond, 2005; Hervio, Coombs, Bergstrom, Trivedi, Corey & Madison, 2000). Penalty: 0.1; sort order: average score. Similarity score: 100; specificity, >95%. *b*. Effects of plasmin on αβγΔ131-138 + 3A (K168A, R178A, and K179A) channel activity in 1 and 24 hr. n = 18. NS, not significant, * P < 0.05 vs before. *c*. Immunoblotting assays of αβγΔ131-138 and αβγΔ131-138 + 3A channel proteins. These results are a representative blot of three experiments. *d*. Regulation of single point mutants (γK168A, γR178A, and γK179A) derived from αβγΔ131-138 by plasmin. n = 12. NS, not significant vs αβγΔ131-138 + 3A (3A) construct. *e*. Detection of plasmin-cleaved fragments of three single point mutants (from left to right are αβγΔ131-138, αβγΔ131-138+K168A, αβγΔ131-138+R178A, and αβγΔ131-138+K179A). *f*. Cleavage efficacy (cleaved band/uncleaved band). **P < 0.01. n = 3.

Different size of cleaved band of γΔ178-193 mutant by plasmin indicates additional cleavage sites may exist. Based on the prediction (Figure 7a), we constructed a new mutant termed γΔ178-193 + 4A (K126A, R135A, K136A, and R153A). The activity of this mutant was not activated by plasmin (Figure 7b, Table 1). Moreover, the “full-length” proteins of this mutant were increased significantly compared with the γΔ178-193 mutant (Figure 7c). This was further confirmed by the densitometrical ratio of cleaved/full-length proteins (Figure 7d), suggesting that plasmin could cleave multiple sites in the finger domain of γENaC (Figure 7e).

**FIGURE 7.**
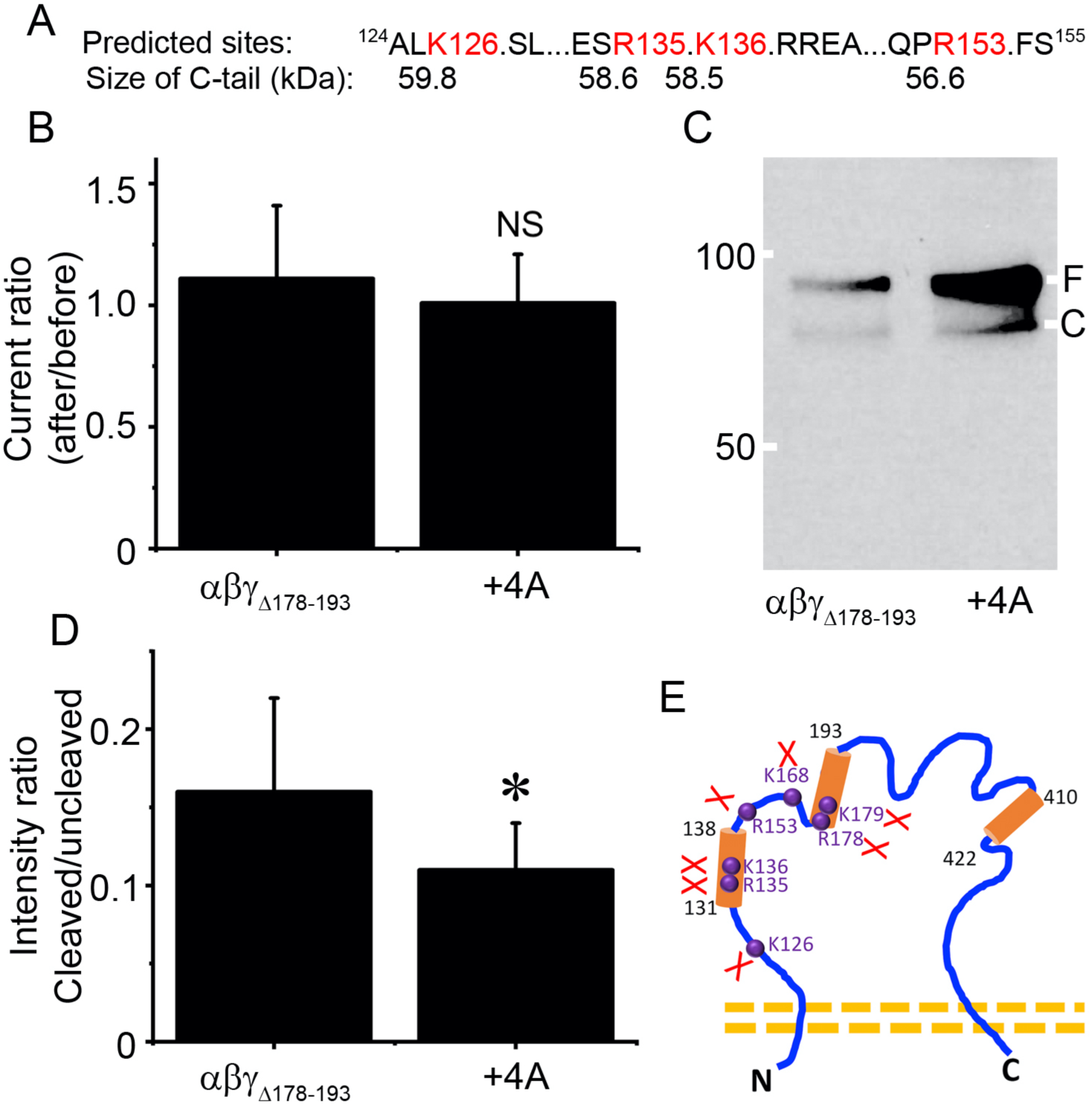
Prediction and validation of putative plasmin cleaved sites preceding γK168. *a. In silico* predicted four potential cleavage sites (K126, R135, K136, and R153) and the peptide size of C-terminal fragments. *b*. Effects of plasmin on the channel activity of αβγΔ178-193 and a mutant combining γΔ178-193 and four predicted sites (+4A) that were replaced with alanine. n = 9 - 14. NS, not significant vs αβγΔ178-193 mutant. *c*. Immunoblotting assays. This blot represents four experiments with similar results. *d*. Contribution of four predicted cleavage sites. *P < 0.05 vs αβγΔ178-193 construct. n = 4. *e*. Summary of newly identified and validated cleavage sites for plasmin in γ ENaC subunit. Three putative proteolytic regions for truncation mutants are marked with cylinders, four identified amino acid residues are labeled in purple font and balls.

**FIGURE 8.**
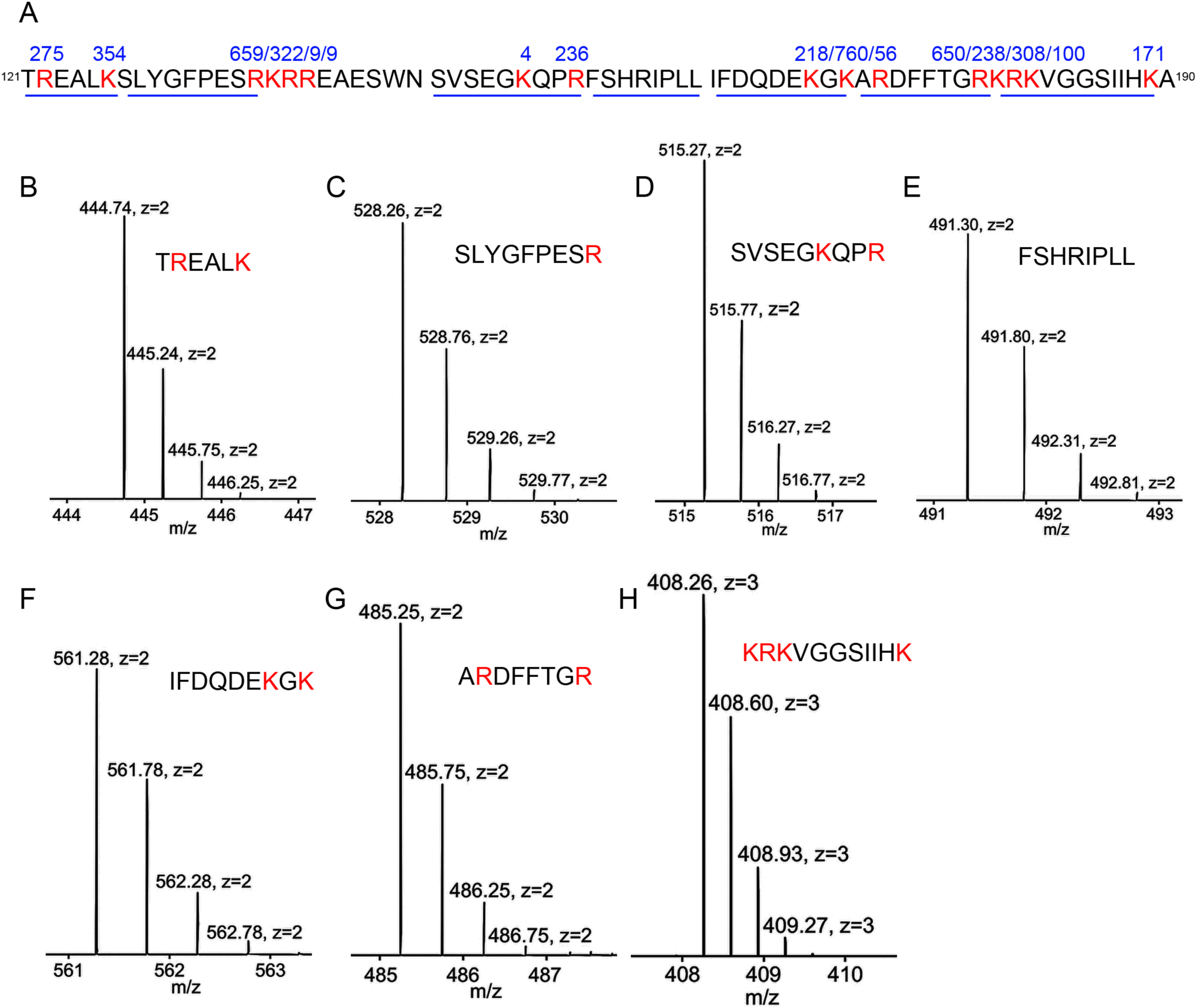
Validation of plasmin cleaved sites by mass spectrometry. *a*. Sequences of three synthesized peptides with identified plasmin cleavage sites (red font). Three peptides with a continuous sequence were separated by two gaps. The numbers above the cleaved sites are the frequency of the fragments with an end of this amino acid residue identified by mass spectrometry. The underlines indicate the sequences of the fragments for panels *b* to *h* from left to right. *b-h*. MS1 spectrum for the representing fragments. The size and charge are labeled for each isotope. Corresponding sequences of each fragment occurs as insets.

### 3.7 Validation of plasmin cleavage sites by mass spectrometry (MS)

To corroborate identified cleavage sites, three synthesized peptides identical to the sequence of γENaC from ^121^T to A^190^ amino acid residues were treated with plasmin and analysed by LC-MS. In addition to confirming the identified critical residues, nine more cleavage sites were detected (Figure 8a and Table 1). The frequency for the fragments with either the R or K amino acid residue as the C-terminal tail or the next amino acid residue as the N-terminal tail was computed, the confirmed 7 cleavage sites identified by mutagenesis and confirmed by MS occurred more than 200 times. In comparison, the frequency of most MS discovered new cleavage sites was less than 100 except R122, K170, R180, and K189 (Figure 8a). Representative MS1 spectrum of seven fragments were shown in Figure 8b-h. In general, 4-5 isotopes with variant intensity/counts for the same fragment were seen. Corresponding MS/MS spectrum for these fragments was included in the Supplementary data (Figures S1-S7).

### 3.8 Identified cleavage sites for plasmin compose two proteolytic centres

To address the hypothesis that the identified 16 sites may be comprise of cleavage centres spatially in 3D structure (Figure 9a), we labeled three deletion mutants in key domain organization scheme (Figure 9b) and GRIP domain (Figure 9c) based on the cryo-EM model (Noreng, Bharadwaj, Posert, Yoshioka & Baconguis, 2018). Apparently, one center was made up of six cleavage sites at the end of α1 helix and another with cleavage sites located in antiparallel P1 and P2 strands. On the other hand, we employed protein docking approach to substantiate this notion and potential interactions between individual amino acid residue. We generated a homology model of human γENaC with the I-TASSER server (Figure S8a)(Noreng, Bharadwaj, Posert, Yoshioka & Baconguis, 2018; Roy, Kucukural & Zhang, 2010). There were three highly accessible clusters of positively charged residues in the GRIP domain of the γ ENaC (Figure S8b). As positive control, Figure S9c-d show interactions between plasmin and two arginine residues (R17 and R19) of textilinin-1 (pdbID: 3uir). Using the interacting residues in plasmin, we found the best docking result between 178RKRK in γ ENaC and D735 and G764 of plasmin (Figure S8e-f). Further, interactions between 135RKRR of γ ENaC and plasmin (Figure S8g-h). Together, two spatial centres for the cleavage by plasmin are identified in the vicinity of α1 terminal and antiparallel P1/P2 strands.

**FIGURE 9.**
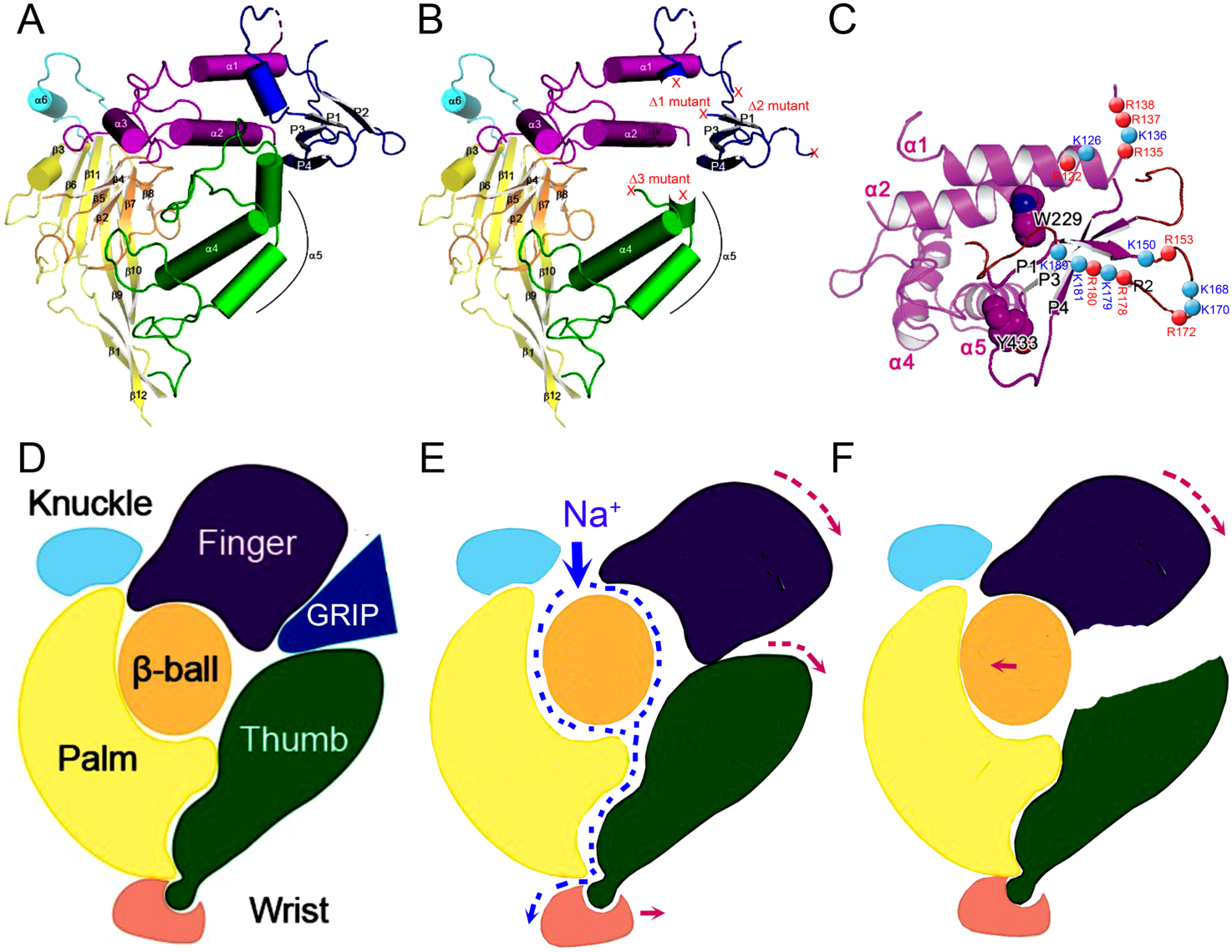
Location of deletion and single or multiple point mutants in 3D model and gating mechanism. *a*. 3D structure of truncated human ENaC subunits adapted from Noreng S 2018. *b*. Location of deleted regions (Δ1, Δ 2, and Δ 3) for both α and γ ENaC subunits. *c*. Positions of identified amino acid residues for plasmin cleavage in human γ subunit. *d-f*. Schematic mechanism for plasmin to cleave and gate ENaC protein complexes. Plasmin cleaves full-length ENaC (*d*) to remove the GRIP domain (*e*). The finger domain will subsequently fall down to hit the thumb to increase the width of channel pore. Na^+^ ions will go through channel pore faster as shown by increased currents. However, for some mutants, for example, Δ3 deletion mutants of α and γ subunits, the link between the finger and the thumb disappears so that downward moved finger cannot reach the thumb to broaden channel pore (*f*).

### 3.9 G protein pathway may be involved

ENaC was regulated by G proteins, which were downstream molecules of the plasmin signal pathway (Greenlee et al., 2013). Because the effects of plasmin lasted up to 24 h, plasmin could regulate ENaC via activation of the G protein signal. To exclude this possibility, pertussis toxin was used to specifically inactivate Gi/o activity. Activation of ENaC currents by uPA and plasmin was not affected significantly (Figure S9a). However, the fold of plasmin-activated current was significantly reduced compared with controls and uPA groups (Figure S9b). These data suggest that plasmin but not uPA could activate ENaC by activating the G protein signal pathway as a long-term effect.

## 4 DISCUSSION

Given the pharmaceutical features of plasmin and depressed fibrinolytic activity in ARDS, we set out to address the hypothesis that plasmin could improve AFC by cleaving ENaC proteolytically. We hereby provide preclinical evidence showing that intratracheally administered plasmin augmented AFC in both normal and acid aspiration injured lungs. For the first time, we found that plasmin catalysed both full-length and furin-trimmed human γENaC proteins. Plasmin may be a newly identified pharmaceutical intervention for removing oedema fluid from the air spaces in ARDS.

We identified novel cleavage sites for plasmin beyond three putative regions described previously (Kleyman, Carattino & Hughey, 2009; Planes & Caughey, 2007; Rossier & Stutts, 2009) (Table 1). The GRIP domain (gating relief of inhibition by proteolysis) is comprised of four β strands (refereed as P1-4 segments) (Figure 9a & d)(Noreng, Bharadwaj, Posert, Yoshioka & Baconguis, 2018). Plasmin cleaved 16 sites, at least two of them are out of this GRIP domain but within the α1 helix of the finger. The antiparallel arrangement of P1 strand, P2 strand, and the α1 helix place all of plasmin cleavage sites in close proximity to facilitate proteolytic efficiency of plasmin by forming two proteolytic centres. Plasmin is the most powerful serine protease and shares cleavage sites with prostasin, trypsin I and IV, uPA, elastases, chymotrypsin, and furin.

One of our novel observations is that long-term exposure of human ENaC proteins to plasmin leads to the cleavage of full-length γ ENaC. The current concept is that serine proteases could not cleave full-length ENaC but furin-cut C-terminal fragment potentially due to the uncovery of hiding catalytic triad by furin for chymotrypsin and others (Kleyman, Carattino & Hughey, 2009; Rossier & Stutts, 2009). Similar to furin, plasmin may cleave furin sites and expose additional proteolytic sites for itself. Our findings that long-term exposure to plasmin hydrolyses full-length ENaC proteins shifted the concept that only furin-cleaved ENaC proteins are accessible for external proteinases. This may explain why plasmin exhibited more potency to activate ENaC compared with tc-uPA. Additionally, plasmin but not tc-uPA activated the G-protein pathway may facilitate ENaC activation too.

The differences between immunoblotting and functional assays in the third deletion mutants, (i.e., αΔ432-444 and γΔ410-422) could be due to the disassociation of the finger and the thumb. Removal of the GRIP domain by proteolysis results in a downward movement of the finger, subsequently hitting the thumb to enlarge the channel pore and gate. Eventually, the channel activity is up regulated maximally (Figure 9e). Even though cleavage of the GRIP domain by plasmin, as shown on Western blots, still leads to fall down of the finger, truncated thumb of these deletion mutants could not be hit by the fallen finger, which would not alter the size of channel pore and function. In addition, deletion of these putative proteolytic regions may alter the formation of channel pore. This hypothesis is supported by the first 3D model of human ENaC proteins (Noreng, Bharadwaj, Posert, Yoshioka & Baconguis, 2018).

Our *in silico* prediction excludes the cleavage of human α, δ, and β subunits by plasmin. To date, only α and γ ENaC subunits are predominately cleaved by tested proteases. Although δ subunit affects the cleavage of these two subunits, cleavage of δENaC has not been reported. The cleavage of βENaC by serine proteases is rare and weak (Garcia-Caballero, Dang, He & Stutts, 2008; Jovov, Berdiev, Fuller, Ji & Benos, 2002). Based on our *in silico* prediction and functional results, the possibility for plasmin to cleave β subunit is much lesser compared with γ counterpart.

The greater real size of C-terminal fragments on blots compared with the prediction by the server could be due to post-translational modifications. Similar to our observations, up to 110 kDa of αβγENaC proteins was described, which was even larger than what we detected. Recently, we executed mass spectrometry for the two bands of δENaC on western blots. Both the band with a predicted size (75 kDa) and a large band (110 kDa) were confirmed to have δENaC sequence (Zhao et al., 2019). Therefore, we deduce that cleavage of ENaC by plasmin could facilitate post-translational modifications.

We could not completely rule out the involvement of protease-activated receptor-1, -2 and -4 (PAR) to fluid resolution in human and mouse lungs (Bock et al., 2015; Carmo et al., 2014; Mannaioni et al., 2008; Quinton, Kim, Derian, Jin & Kunapuli, 2004). However, activation of lung epithelial PAR isoforms by plasmin have not been reported.

Under physiological conditions, bronchoalveolar fluid plasmin (1.5 μmol·L^-1^) may activate ENaC indirectly through cleavage of sc-uPA to produce active tc-uPA. However, this possibility could be ruled out in diseased lungs. In the bronchoalveolar lavage fluid of ARDS, antiplasmin molecules, including antithrombin, α1-antitrypsin, α2-antiplasmin, and α2-macroglobulin are markedly elevated. For example, α1-antitrypsin concentration increased 60-fold in diseased lungs (Wewers, Herzyk & Gadek, 1988). These elevated plasmin inhibitors bound to plasmin overwhelmingly if any to cage its enzymatic activity. On the other hand, increased urokinase inhibitors, PAI-1 would form complexes with uPA and eliminate physical interactions between uPA and plasmin. Therefore, a dose above physiological concentration is needed for treating arterial occlusive diseases and macular oedema.

In summary, this study for the first time demonstrates the beneficial effects of plasmin on alveolar fluid clearance and provides novel mechanisms for how the most potent plasmin to cleave multiple sites of human ENaC.

## Supporting information

all figures

## ACKNOWLEDGEMENTS

This study was supported by NIH grants HL87017, HL095435, HL134828, AHA14GRNT20130034, NSFC 81670010, and AHA16GRNT30780002. The authors thank Ms. Yun Jiang (University of Texas Health Science Centre at Tyler) and Dr. Steven Idell (University of Texas Health Science Centre at Tyler) for their superb technical support and thoughtful discussion. The authors acknowledge Dr. Andrew Lemoff and Dr. Xuemei Luo of the UTSW Proteomics Core for their professional assistance with mass spectrometry analysis.

## Abbreviations

ENaC: epithelial sodium channels;
ARDS: acute respiratory distress syndrome;
PAI-1: plasminogen activator inhibitor-1;
AFC: alveolar fluid clearance;
uPA: urokinase plasminogen activator;
AT2: alveolar type II epithelial cells;
MS: mass spectrometry;
GRIP: gating relief of inhibition by proteolysis;
AS: amiloride sensitive.

## AUTHOR CONTRIBUTIONS

RZ Zhao, D Bhattarai, and HL Ji expressed ENaC in oocytes, performed voltage clamp recordings, analysed AFC in mice, and analysed results. RZ Zhao carried out mutagenesis, the immunoblotting assays, and sample preparation for mass spectrometry. HG Nie performed *ex vivo* AFC in human lungs. G Ali performed bioelectrical assays in mouse AT2 monolayers. Y Chang performed homology modelling and protein-protein docking. HL Ji and MA Matthay were responsible for experimental design, data analysis, result assembly, and manuscript preparation.

## DECLARATION OF TRANSPARENCY AND SCIENTIFIC RIGOUR

This Declaration acknowledges that this paper adheres to the principles for transparent reporting and scientific rigour of preclinical research as stated in the BJP guidelines for Design & Analysis, Immunoblotting and Immunochemistry, and Animal Experimentation, and as recommended by funding agencies, publishers and other organisations engaged with supporting research.

