## Supplementary material for "Plasmin improves oedematous blood-gas barrier by cleaving epithelial sodium channels": all figures

[1]Department of Cellular and Molecular Biology, University of Texas Health Science Center at Tyler, Tyler, Texas 75708, USA; [2]College of Basic Medical Science, China Medical University, Shenyang, Liaoning 110001, China; [3]Barrow Neurological Institute, 350 West Thomas Road, Phoenix, Arizona 85013-4409, USA; [4]Department of Physiological Sciences, Eastern Virginia Medical School, Norfolk, Virginia 23501, USA; [5]Department of Anesthesia and Medicine, University of California San Francisco, San Francisco, California, USA; [6]Texas Lung Injury Institute, University of Texas Health Science Center at Tyler, Tyler, Texas 75708, USA

**FIGURE S1-7**. Online supplementary figures are the representative MS/MS spectrum for the peptide fragments in Figure 8.

Figure S1

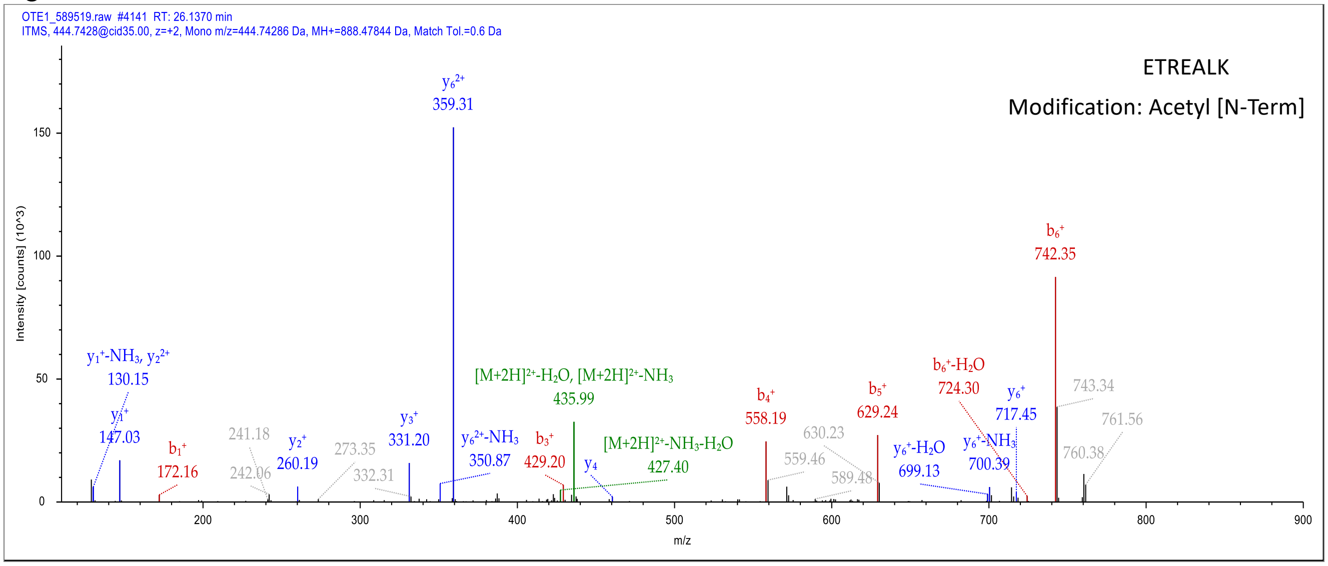


Figure S2

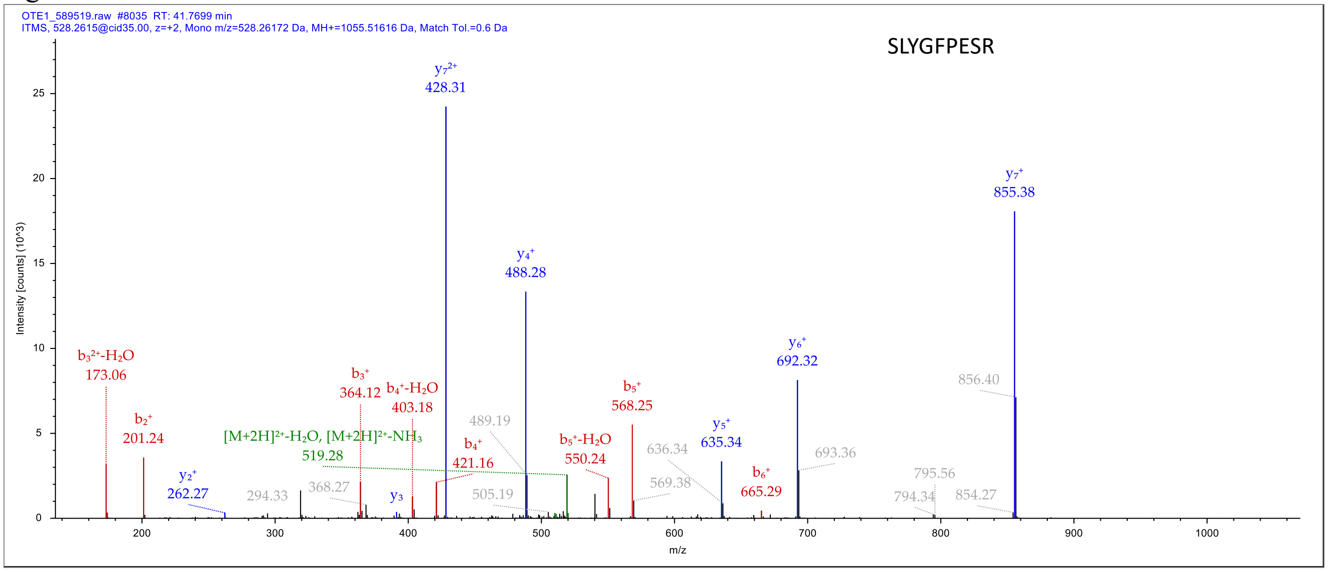

### Figure S3

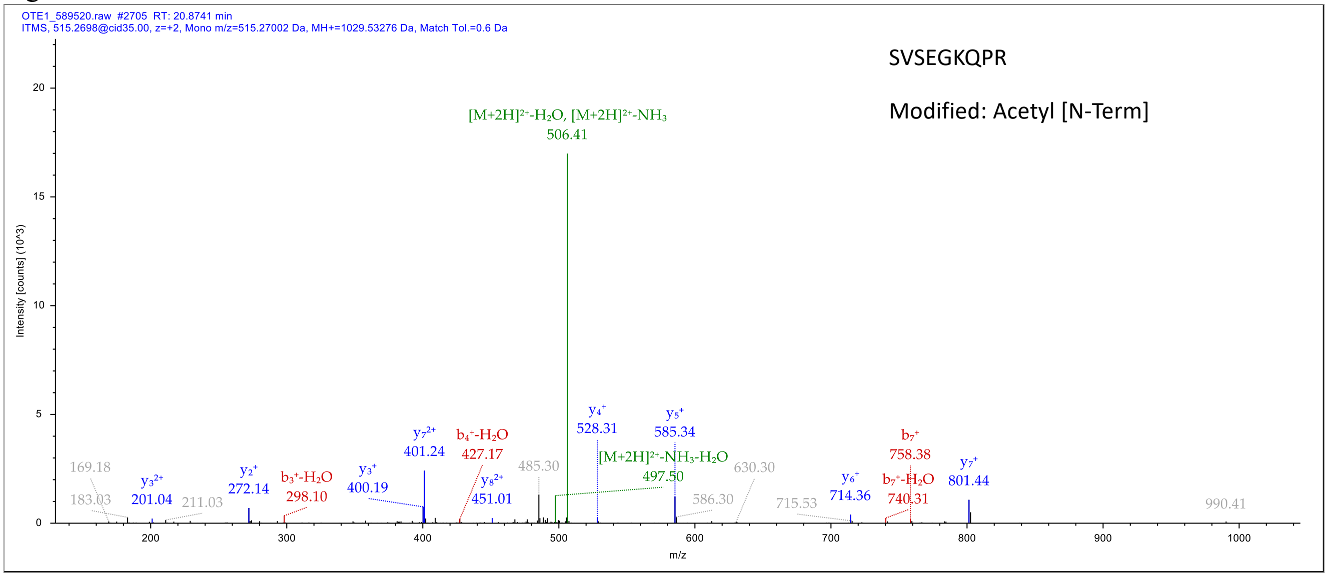


### Figure S4

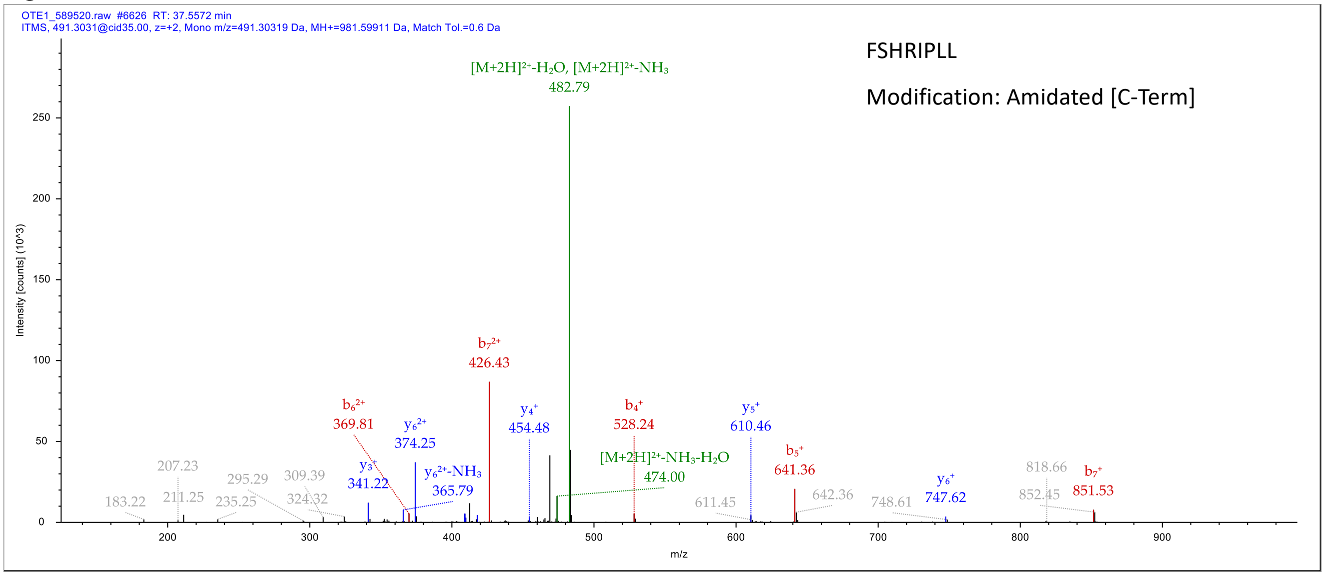

Figure S5

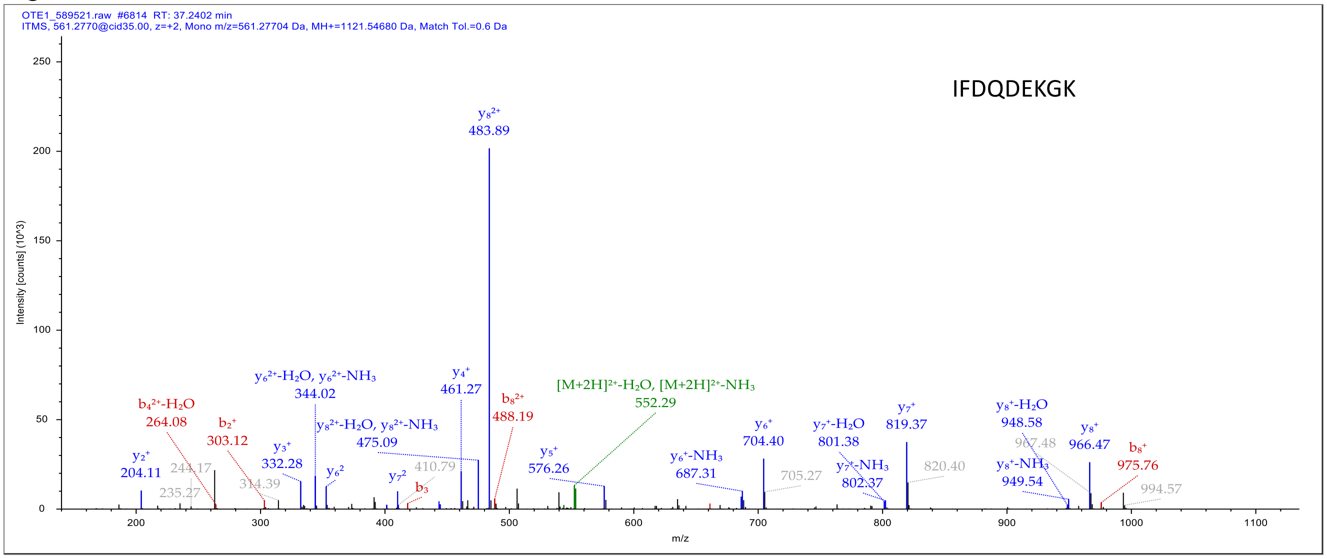


Figure S6

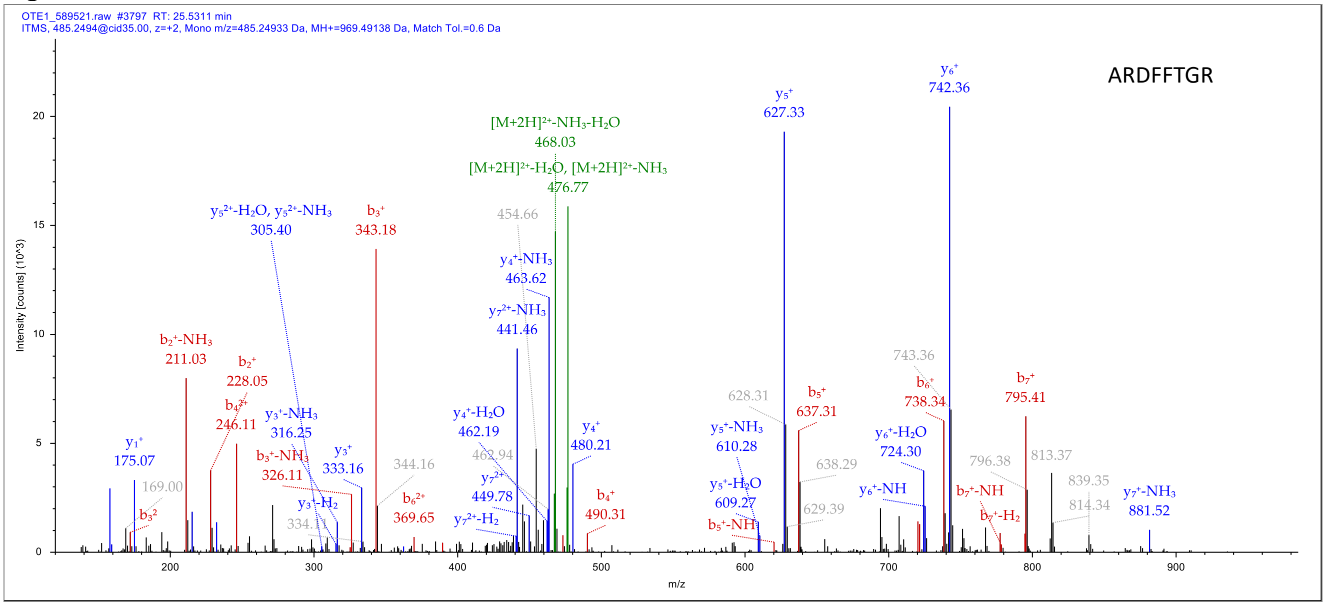

Figure S7

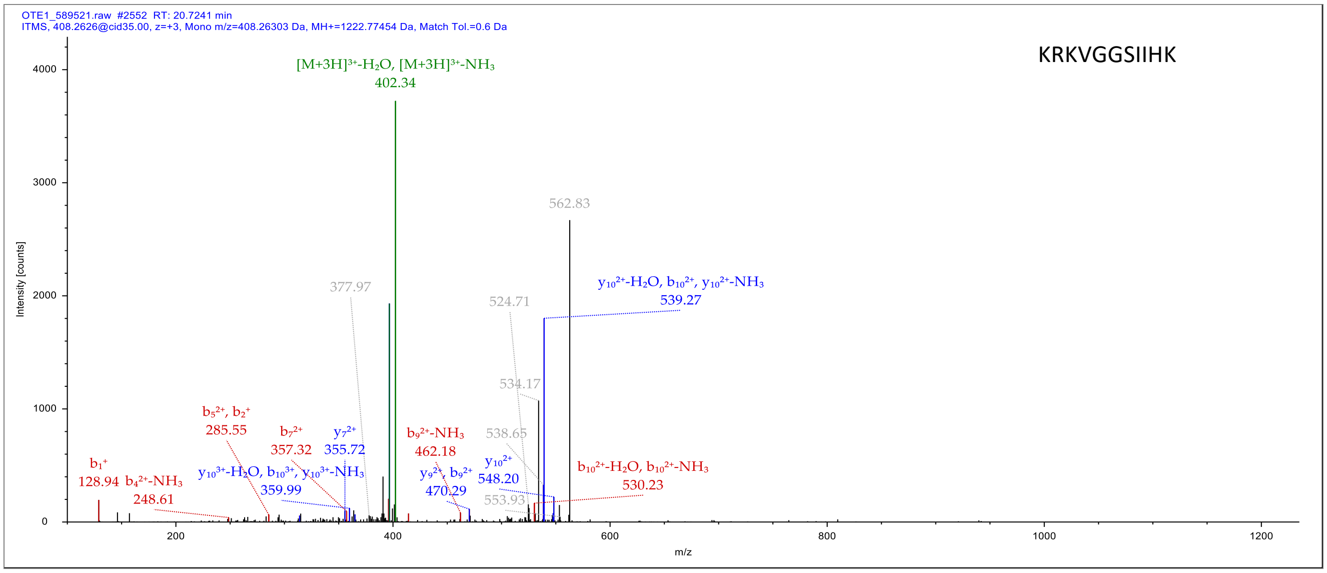

OTE1_589521.raw #2552 RT: 20.7241 min
ITMS, 408.2626@cid35.00, z=+3, Mono m/z=408.26303 Da, MH+=1222.77454 Da, Match Tol.=0.6 Da
KRKVGGSIIHK
Intensity [counts]
m/z
[M+3H]3+-H2O, [M+3H]3+-NH3
402.34
562.83
377.97
524.71
y10 2+-H2O, b10 2+, y10 2+-NH3
539.27
534.17
b5 2+, b2+
285.55
b7 2+
357.32
y7 2+
355.72
b9 2+-NH3
462.18
538.65
b1+
128.94
b4 2+-NH3
248.61
y10 3+-H2O, b10 3+, y10 3+-NH3
359.99
y9 2+, b9 2+
470.29
y10 2+
548.20
553.93
b10 2+-H2O, b10 2+-NH3
530.23

Figure S8. Homology model and docking results for plasmin cleavage centres. *a*. 3D model of the human γ ENaC generated by I-TASSER. *b*. The GRIP domain (aa114-239) of the human γ ENaC model with three positively charged residue clusters facing outward and highly accessible. Four P strands are composed of following amino acids: P1, aa160-166; P2, aa182-190; P3, aa199-208; and P4, aa213-219. Three cleavage centres were highlighted. *c*. The interaction of plasmin (chain A with surface presentation) with textilinin-1 (chain C) from the original crystal structure 3uir highlighted with two arginine residues in the substrate interacting with plasmin. *d.* Detailed interactions between two arginine residues and plasmin: Arg17 interacting with Asp735, Ser741, and Gly764; Arg19 interacting with Glu687 and Phe587. We used this as a docking guide as attractors (Asp735 and Gly764). The colour of the substrate in *d, f*, and *h* is set yellow for clarity. *e*. Docking result with 178RKRK in the γ ENaC as attractors to the plasmin, showing that two arginine residues Arg178 and Arg180 docked to the plasmin to the similar positions in *c*. *f.* The detailed interactions of Arg178 and Arg180 with the plasmin: Arg178 interacting with Asp735 and other residues (Tyr774, Gln738); Arg180 interacting with Glu623, His586. *g*. Docking result with 135RKRR in the γ ENaC as attractor to the plasmin active centre also resulted in two arginine residues (Arg137 and Arg135) interacting with the plasmin active centre. *h*. The detailed interactions of Arg137 and Arg135 with the plasmin residues as shown.

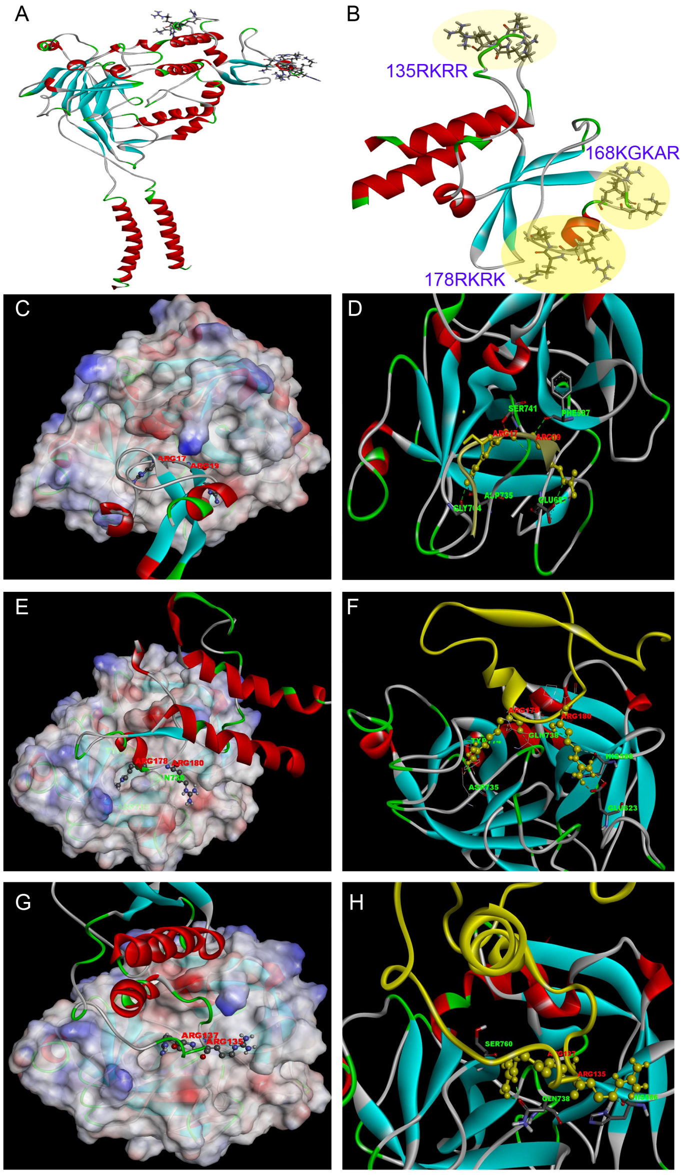

Figure S9. Plasmin regulates ENaC activity via G protein. *a*. ENaC activity in the absence (Control) of two-chain urokinase plasminogen activator (tc-uPA, 10 μg·ml$^{-1}$, 30 min at room temperature) and plasmin (10 μg·ml$^{-}$1, 30 min at room temperature) and presence of protease and Gi/o protein antagonist, pertussis toxin (PTX, 2 μg·ml$^{-1}$, 24 hr prior to addition of proteases at 20 ºC). n = 12. NS, not significant; *P < 0.05 and **P < 0.01, respectively, correspondingly vs those before application of proteases. *b*. Normalized ENaC activity. ** P < 0.01 vs that in the absence of PTX. n = 12.

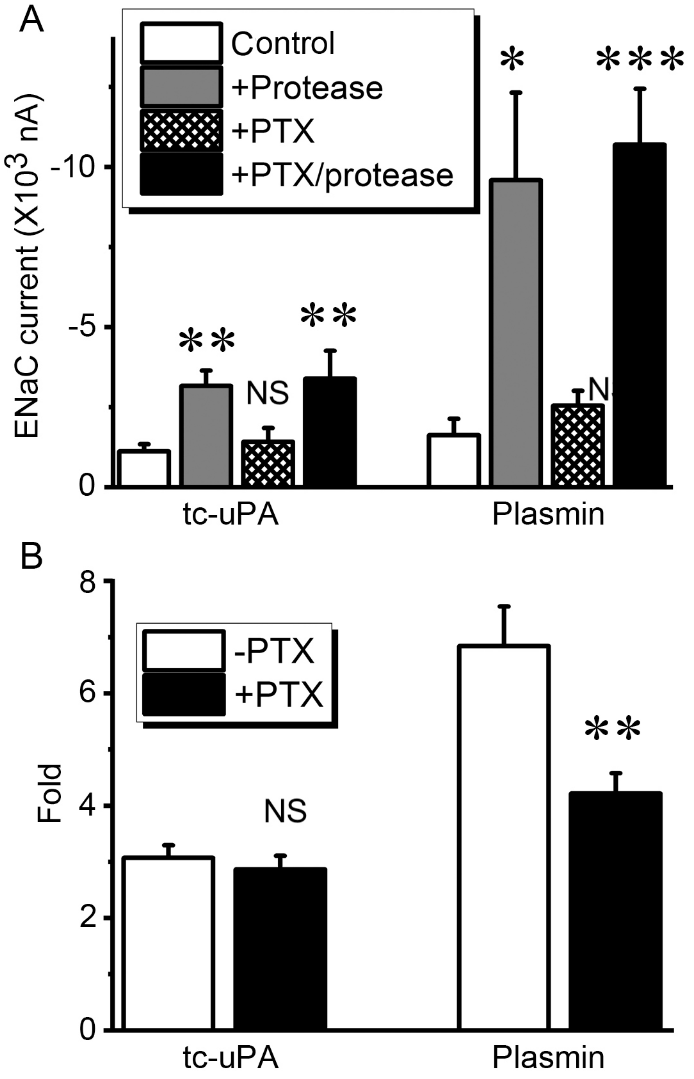
